# Microglia and macrophages alterations in the CNS during acute SIV infection: a single-cell analysis in rhesus macaques

**DOI:** 10.1101/2024.04.04.588047

**Authors:** Xiaoke Xu, Meng Niu, Benjamin G. Lamberty, Katy Emanuel, Andrew J. Trease, Mehnaz Tabassum, Jeffrey D. Lifson, Howard S. Fox

## Abstract

Human Immunodeficiency Virus (HIV) is widely acknowledged for its profound impact on the immune system. Although HIV primarily affects peripheral CD4 T cells, its influence on the central nervous system (CNS) cannot be overlooked. Within the brain, microglia and CNS-associated macrophages (CAMs) serve as the primary targets for HIV, as well as for the simian immunodeficiency virus (SIV) in nonhuman primates. This infection can lead to neurological effects and the establishment of a viral reservoir. Given the gaps in our understanding of how these cells respond *in vivo* to acute CNS infection, we conducted single-cell RNA sequencing (scRNA-seq) on myeloid cells from the brains of three rhesus macaques 12-days after SIV infection, along with three uninfected controls. Our analysis revealed six distinct microglial clusters including homeostatic microglia, preactivated microglia, and activated microglia expressing high levels of inflammatory and disease-related molecules. In response to acute SIV infection, the population of homeostatic and preactivated microglia decreased, while the activated and disease-related microglia increased. All microglial clusters exhibited upregulation of MHC class I molecules and interferon-related genes, indicating their crucial roles in defending against SIV during the acute phase. All microglia clusters also upregulated genes linked to cellular senescence. Additionally, we identified two distinct CAM populations: CD14^low^CD16^hi^ and CD14^hi^CD16^low^ CAMs. Interestingly, during acute SIV infection, the dominant CAM population changed to one with an inflammatory phenotype. Notably, specific upregulated genes within one microglia and one macrophage cluster were associated with neurodegenerative pathways, suggesting potential links to neurocognitive disorders. This research sheds light on the intricate interactions between viral infection, innate immune responses, and the CNS, providing valuable insights for future investigations.

**AUTHOR SUMMARY:** 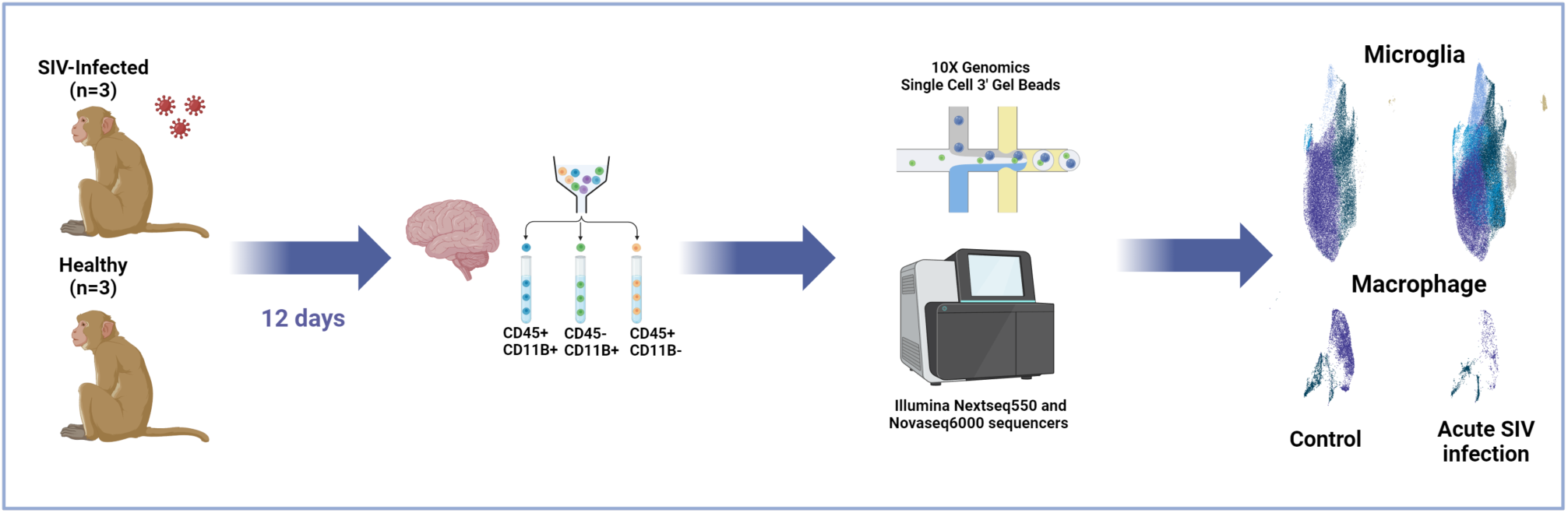

HIV’s entry into the central nervous system (CNS) can lead to neurological dysfunction, including HIV-associated neurocognitive disorders (HAND), and the establishment of a viral reservoir. While microglia and CNS-associated macrophages (CAMs) are the primary targets of HIV in the CNS, their responses during acute HIV infection remain poorly defined. To address this, we employed the scRNA-seq technique to study microglial and CAM populations in rhesus macaques during acute SIV infection. By identifying signature genes associated with different phenotypes and mapping them to various biological and pathological pathways, we discovered two myeloid cell clusters strongly linked to neurodegenerative disorders. Additionally, other clusters were associated with inflammatory pathways, suggesting varying degrees of activation among different myeloid cell populations in the brain, possibly mediated by distinct signaling pathways. All microglia clusters developed signs of the cellular senescence pathway. These findings shed light on the immunological and pathological effects of different myeloid phenotypes in the brain during acute SIV infection, providing valuable insights for future therapeutic strategies targeting this critical stage and aiming to eliminate the viral reservoir.

## INTRODUCTION

The human immunodeficiency virus (HIV) is an enveloped retrovirus that contains two copies of a single-stranded RNA genome, which can cause acquired immunodeficiency syndrome (AIDS) by significantly impairing the immune system. HIV remains a global health challenge with profound implications for individuals, communities, and societies. The estimated number of people with HIV (PWH) is 36.7 million worldwide as of 2016, and according to the latest epidemiology study in 2020, PWH comprise approximately 1.2 million in the United States.(^1^) Similar to HIV in genomic, structural, and virologic perspectives, the simian immunodeficiency virus (SIV) also belongs to the primate retrovirus family. *In vivo*, both viruses cause persistent infection. Infection of nonhuman primates (NHPs) by SIV mimics many key aspects of HIV infection in humans, including immunodeficiency, opportunistic infections, and CNS infection which can be associated with neurological impairments.(^2–5^)

The development of HIV infection can be classified into three stages, acute HIV infection, chronic HIV infection, and if untreated, eventually AIDS.(^6^) The acute infection period is defined as the stage immediately after HIV infection and before the development of antibodies to HIV, during which the virus rapidly multiplies and spreads throughout the body.(^7^) Sexually-mediated HIV transmission generally starts with mucosal CD4+ T cells and Langerhans cells,(^8^) and then travels to gut-associated lymphoid tissue (GALT). Intravenous infection-mediated HIV transmission leads to initial infection of CD4+ T cells in lymph nodes, the spleen, and GALT.(^9, 10^) Lymphoid tissue and other organ macrophages can also be infected. HIV-infected cells can produce large amounts of virus to infect additional target cells and can also migrate and carry the virus to other tissues and organs including the central nervous system (CNS). The earliest post-infection time for detecting HIV/SIV RNA in the CNS (brain or cerebrospinal fluid) ranges from 4 days to 10 days.(^11–13^) Like the deteriorative effect of HIV in the periphery, the inflammatory events and neurotoxicity elicited by the HIV/SIV can damage neurons as well as other supportive cells in the brain which can eventually lead to HIV-associated neurocognitive disorders (HAND). While the extensive use of antiretroviral therapy (ART) has significantly reduced the occurrence of dementia, the most severe type of HAND, the overall prevalence of HAND still hovers around 50%.(^14–17^)

In CNS HIV/SIV infection, CNS-associated macrophages (CAMs) and microglia are thought to play a central role in defending against the invading pathogen and trigger neuroinflammation.(^18, 19^) Responding to the virus and/or virally-infected cells that enter the brain as the initial innate immune response, they are activated and can release numerous proinflammatory cytokines including interferons, IL-6, IL-1β, and TNF-α to contribute to the control and clearance of the virus or infected cells from the CNS.(^11^) However, macrophages and microglia can also contribute to the pathological events of HIV/SIV infection. In acute SIV/HIV infection, infected blood monocytes represent another cell type other than CD4+ T cell that has been proposed to seed the virus in the CNS. Infection of rhesus monkeys with SIV indeed results in an increase in monocytes trafficking to the brain.(^20^) Once trafficking monocytes enter the brain, they can further differentiate to CAMs, and under experimental conditions (such as depletion of CD8+ cells) can lead to rapid development of SIV encephalitis.(^21^) On the other hand, blocking of an integrin (α4), which is highly expressed on monocytes, by natalizumab was found to profoundly hinder CNS infection and ameliorate neuronal injury.(^22^) However integrin(α4) is also expressed on CD4+ T cells, thus attributing the effect to monocytes is uncertain. Although microglia, which are the CNS-specific resident myeloid cells, do not participate in bringing the HIV/SIV to the CNS, they are actively involved in neuronal damage once they are infected and/or activated. (^14, 23, 24^) In addition, infected CAMs and/or microglia are thought to make up a viral reservoir in the brain under suppressive ART treatment, complicating efforts for an HIV cure.

Although the general responses of microglia and CAMs to acute SIV infection has been widely studied by using bulk assays, our understanding of different microglial or CAMs phenotypes have is minimal for this important period in which virus enters the brain. In this study, we performed high-throughput single-cell RNA sequencing (scRNA-seq) on microglia and CAMs from the brains of rhesus macaques during acute SIV infection as well as in control uninfected animals to address the limitations of bulk assays and investigated the different effects of acute SIV infection on varied myeloid phenotypes. We identified homeostatic microglia, preactivated microglia, activated microglia, and two phenotypes of CAMs in the brains. We further characterized these subsets of cells by comparing their transcriptomic profiles in the uninfected and acute-infected conditions. Different responses were evoked in different microglial and macrophage phenotypes in acute SIV infection. Interestingly, there were two activated cell clusters that were found to be closely associated with neurological disorders. Finally, although we did not observe modulation of the expression of the anti-apoptotic molecule BCL-2 (upregulation of BCL-2 is one of the mechanisms for promoting the survival of infected cells) (^25, 26^) by SIV in microglia and CAMs, we found another anti-apoptotic molecule, CD5L was highly expressed in infected microglia which might be a novel potential pathway to elucidate the reservoir establishment in the CNS.

## RESULTS

### The constitution of the brain’s myeloid cell population was altered by acute SIV infection

To examine the effect of acute SIV infection on brain myeloid cells, we inoculated three rhesus macaques with SIV_mac_251, and processed the brains at 12-days post-inoculation to enrich resident and infiltrating immune cells for scRNA-seq. Cells from three uninfected animals were similarly analyzed. In the infected animals, high viral loads were found in the plasma at this stage, and productive brain infection (high viral RNA/DNA ratios) was found **(Figure S1, Table S1)**. Uniform Manifold Approximation and Projection (UMAP) visualization of the scRNA-seq data revealed that the transcriptional patterns of the cells purified from the brains of the six rhesus macaques (three uninfected and three acute SIV-infected) grouped in five distinct regions. Graph-based clustering revealed that two of those regions contained distinct clusters, whereas in the other three existed additional subclusters of cells, totaling fifteen separate clusters **(Figure 1A)**.

**Figure 1.**
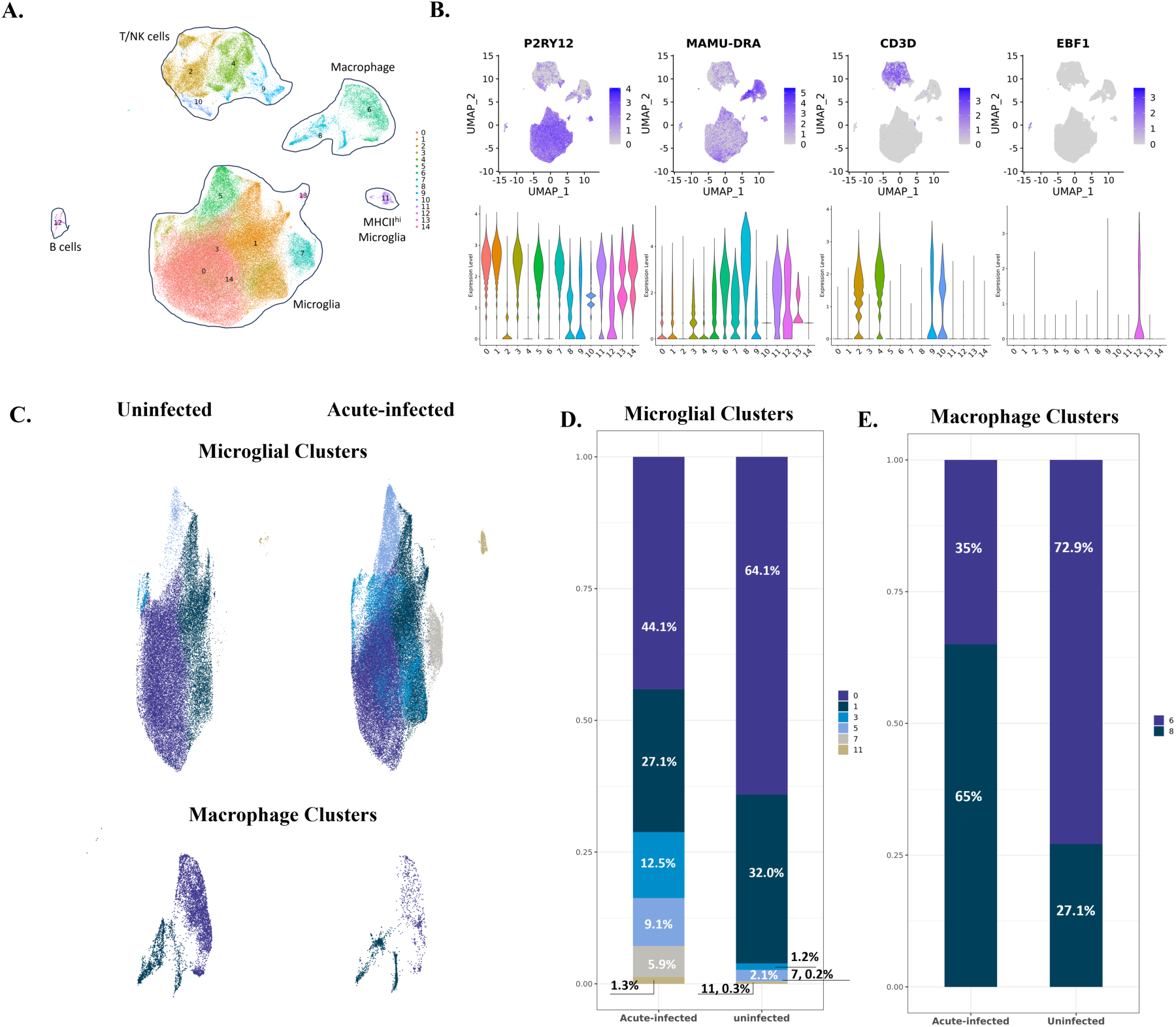
The identified cell clusters in the brains of uninfected and acutely infected rhesus macaques, and the population changes of myeloid cells. **(A)** Fifteen cell clusters (cluster 0 to cluster 14) were identified by graph-based clustering and projected in UMAP. Cluster 0, 1,3, 5, 7, 11, 13, 14 belong to microglia; cluster 2, 4, 9, 10 belong to T/NK cells; cluster 6 and 8 belong to macrophage, and cluster 12 belongs to B cells. **(B)** The expression of representative cell markers used to identify microglia (P2RY12), macrophages (MAMU-DRA), T cells (CD3D) and B cells (EBF1). **(C)** The comparison of myeloid cells’ compositions in the brain between uninfected animals and animals with acute SIV infection. The upper UMAPs include the microglial cluster 0, 1, 3, 5, 7, 11 and the lower UMAPs include macrophage cluster 6 and 8. **(D)** Percentage of each microglial cluster in animals with acute infection and uninfected animals. **(E)** Percentage of each macrophage cluster in animals with acute infection and uninfected animals.

After comparing the expression of different cell markers between those 15 cell clusters **(Figure 1B and Table S2)**, we identified seven microglial clusters with high expression of microglial core genes (P2RY12, GPR34, and CX3CR1) within the main microglia cluster, and one separate cluster of microglia-like cells that also expressed high levels of major histocompatibility complex class II (MHC II) molecules. We also found two macrophage clusters, which had high expression of MHC II and S100 molecules (MAMU-DRA, CD74, S100A6, and S100A4). In addition, we identified four T/NK cell clusters with high expression of the genes for T cell co-receptors, granzymes, and NK cell granule proteins (CD3D, GZMB, and NKG7), and one B cell cluster with high expression of B cell markers (EBF1 and MS4A1). The scope of this study is to investigate the myeloid cells in response to the acute SIV infection, so we only included the myeloid cells, microglia and CNS-associated macrophages (CAMs), for subsequent analyses.

We first assessed whether we could detect cells expressing SIV transcripts. Indeed, in both the microglia and CAM clusters, SIV-infected cells could be identified, in total 0.15% of these myeloid cells (**Table S1**). Given the sensitivity of scRNA-seq, this percentage may be artificially low due to false negatives. In the five regions of brain analyzed for SIV DNA from the three acutely infected monkeys, we found an average of 162 copies of the SIV proviral genome per million cells **(Figure S1, Table S1)**. If one approximates brain macrophages and microglia as comprising 10% of CNS cells, this would imply that 0.16% of these cells are infected, quite similar to our finding of 0.15% of these cells with SIV transcripts. Thus, at least during the acute infection stage, it is likely that the level of expression from the viral genome is sufficient to be recognized in scRNA-seq experiments. Given the similarity in the proportion of cells with SIV DNA and SIV transcripts, it is likely that most infected cells are expressing SIV transcripts, consistent with the high SIV RNA to DNA ratio (**Figure S1, Table S1**).

We next examined whether the infected myeloid cells overall exhibited differences in their gene expression patterns. Other than SIV itself, only two genes (LOC100426197, a class I histocompatibility gene; and LOC693820, the 40S ribosomal protein S29) reached statistical significance for differential expression between the infected myeloid cells and uninfected ones in the acutely infected animals (**Table S3**). This is likely due to activation of even the uninfected cells in the infected animals, as comparing the SIV-positive cells to all the myeloid cells in the uninfected animals yielded 226 DEGs (**Table S3**). Confirming the effect of the acute infection on uninfected cells, when comparing these bystander uninfected cells from the acutely infected animals to those in the uninfected animals, 428 DEGs were identified (**Table S3**). 182 DEGs were in common between the last two comparisons (**Figure S2**).

To better to assess how acute infection alters the myeloid populations, we then separately compared the different microglia and macrophage clusters between the acutely infected animals and the control uninfected animals. Cluster 13 and cluster 14 were excluded for their low cell population (cluster 13 has 273 cells and cluster 14 has 177 cells) and fewer identified markers other than microglial core genes **(Table S2)**. Therefore, we did downstream analyses on the six microglial clusters (0: Micro-0, 1: Micro-1, 3: Micro-3, 5: Micro-5, 7: Micro-7, 11: Micro-11) and two macrophage clusters (6: Macro-6 and 8: Macro-8).

As described briefly above, the initial identification of cell phenotypes revealed that Micro-11 had a high expression of microglial core genes, but from the UMAP **(Figure 1A)**, it did not aggregate with other microglial clusters. In addition, the cells in Micro-11 also had high expression of MHC class II molecules **(Figure 2A)**. Even though MHC class II molecules typically are not highly expressed on the microglia cells compared with CAMs,(^18^) some of the microglial cells in white matter or under pathological conditions might have higher expression.(^27–29^) Thus we included this cluster in the analyses for microglia instead of CAM.

**Figure 2.**
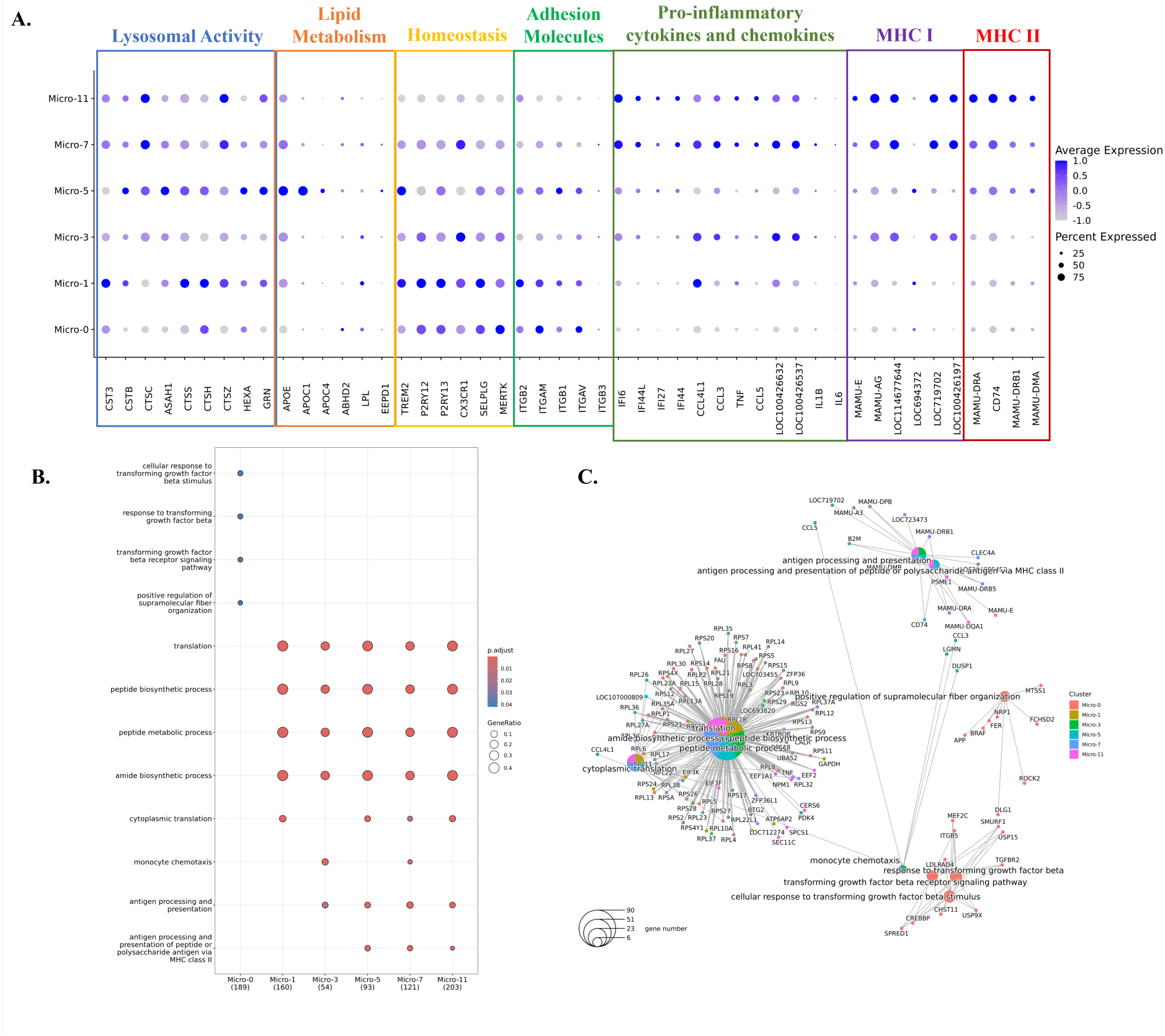
Characterizations of the identified microglial clusters. **(A)** Expression of selected differentially expressed genes (DEGs) in six microglial clusters. The DEGs of each microglial cluster were found by using Wilcoxon rank sum test for comparison. The genes shown were categorized by the functions related to lysosomal activity, lipid metabolism, homeostasis, adhesion molecules, proinflammatory cytokines and chemokines, MHC class I molecules and MHC class II molecules. The data was scaled for plotting. LOC100426537: C-C motif chemokine 3-like. LOC100426632: C-C motif chemokine 4. **(B)** The characteristic biological processes in each microglial cluster by Gene Ontology (GO) pathway analysis. **(C)** The networks between DEGs and the biological processes in each microglial cluster.

Comparing the microglial constituents between the uninfected and infected animals **(Figure 1C and Figure 1D)**, we found that Micro-0 and Micro-1 decreased their proportions under acute SIV infection, which was compensated by the increased proportions of microglial cells from other four clusters. Furthermore, this increase in acute infection was more remarkable for Micro-3 and Micro-7, each increasing at least 10-fold, suggesting those two clusters were highly reactive to the infection. For the macrophage clusters, Macro-6 was the predominant macrophage population comparing to Macro-8 in the brains of uninfected animals (1.9-fold higher), however, this pattern was switched during SIV infection **(Figure 1C and Figure 1E)**. There was a 3.5-fold higher population of Macro-8 over Macro-6 in animals with acute SIV infection, suggesting important roles of Macro-8 in reacting to early viral invasion.

### The increased microglial populations in acute SIV infection were activated microglia but they have different activation patterns and pathways

To better understand the factors behind the changes in microglial populations during acute SIV infection, we further characterized the clusters. We initially compared them regarding the expression of lysosomal proteins, lipid-metabolic proteins, homeostatic molecules, integrins, pro-inflammatory cytokines, chemokines, MHC class I and MHC class II molecules for their important roles in the immune surveillance and maintaining homeostasis of CNS **(Figure 2A)**. These included commonly reported homeostatic genes of microglia include purinergic receptors (P2RY12 and P2RY13), fractalkine receptor (CX3CR1), selectin P (SELPLG), triggering receptor expressed on myeloid cells (TREM2), and tyrosine-protein kinase MER (MERTK). Micro-0 and Micro-1 had higher expression of these genes compared with the other four microglial clusters, indicating they were less likely to be associated with activation and inflammation. However, compared with Micro-0, Micro-1 had enhanced lysosomal activities and higher expression of some inflammatory molecules (APOE, CCL3 and CCL4), suggesting it might be in a slightly more activated, or preactivated state. Micro-3 and Micro-5 both had lower expression of homeostatic genes than Micro-0 and Micro-1, and higher expression of genes associated with microglial activation. Micro-3 highly expressed chemokines CCL3, CCL4 and MHC class I molecules, and Micro-5 highly expressed lysosomal proteins, apolipoproteins (APOE, APOC1 and APOC4) and MHC class II molecules, which suggested that both Micro-3 and Micro-5 might be the activated, or response to infection, potentially disease-related microglia. Given that more inflammatory molecules and cytokines were highly expressed, and the homeostatic genes were rarely expressed in Micro-7 and Micro-11, those two microglial clusters might be in a higher activated state. However, Micro-11 had higher expression of MHC class II molecules with lower expression of homeostatic genes, suggesting it might be the most activated microglial population. Thus, in concert with our findings above the proportion of homeostatic microglia decreases, whereas the proportion of activated microglia increases, during acute SIV infection.

We then implemented GO analyses with the DEGs in each microglial cluster to assess biologic functionalities **(Figure 2B and 2C)**. Micro-0 was identified with the pathways associated with transforming growth factor beta (TGFβ), which further confirmed the homeostatic status of this microglial cluster.(^30^) The other five microglial clusters were all active in the translation and biosynthesis, but Micro-3, Micro-5, Micro-7 and Micro-11 had additional pathways related to microglial activation, such as antigen-presenting ability in MHC class I or MHC class II manner and chemotactic ability. However, the activated microglia clusters appeared specialized in different inflammatory or activation pathways, suggesting the heterogeneity of the activated or pathogenic microglial phenotypes in the brain. In summary, these results indicated that Micro-0 represented homeostatic microglia, Micro-1 might be a preactivated cluster with more activities in protein translation and lysosomal functions, and the other four clusters were the activated microglia with upregulation of different inflammatory molecules and pathways.

### Genes and pathways related to the MHC class I and type I interferon production were upregulated in all microglial clusters responding to the acute SIV infection

In the characterization of the microglial clusters above, we found one homeostatic cluster, one preactivated microglial cluster and four activated clusters. Although the cells from uninfected animals and infected animals were aggregated together for clustering, they still differ in the expression of some genes. Therefore, we identified the genes that were significantly upregulated in each microglial cluster due to acute SIV infection **(Table S4)**. More upregulated genes with high fold-change in acute SIV infection were found in activated clusters Micro-3 and Micro-11, where the average expression of the genes associated with activated microglia were low in uninfected animals **(Figure 3A)**. Therefore, for those two clusters, it might be that only the microglial cells from infected animals were truly activated microglia. For the other activated or preactivated microglial clusters, the cells from uninfected animals did have higher expression of the genes associated with activated microglia compared with the homeostatic cluster Micro-0, and those clusters further upregulated the expression of those activation genes in responses to acute SIV infection.

**Figure 3.**
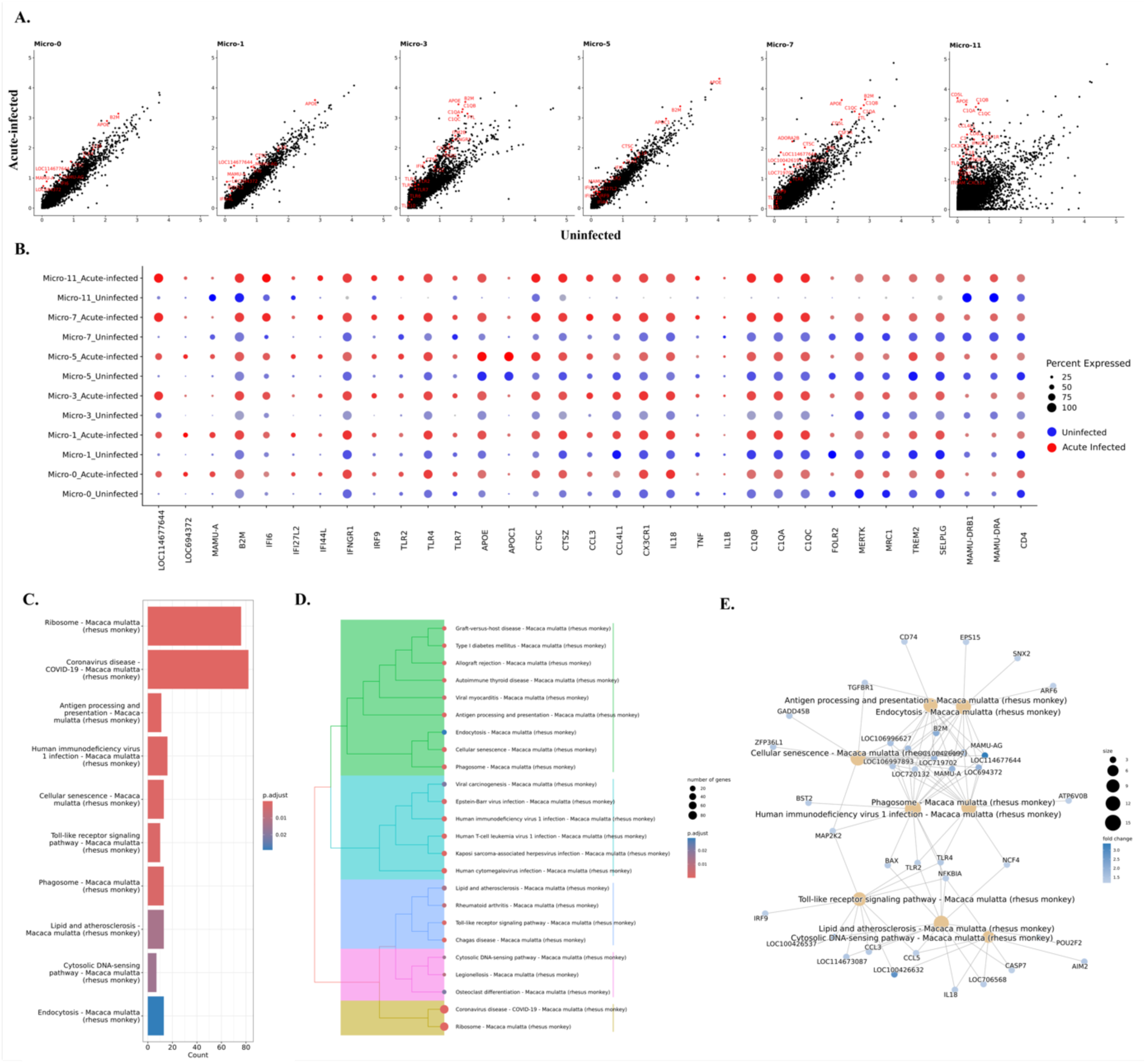
Genes and pathways in microglia that were changed during acute SIV infection. **(A)** The comparison of gene expression levels between the acute-infected animals and uninfected animals for each microglial cluster. Some of the upregulated genes were labeled in red. **(B)** Expression of selected genes that were differentially expressed in uninfected animals (blue dots) and animals with acute infection (red dots) for each microglial cluster. **(C)** Top10 upregulated pathways in microglia from acute infection animals. The upregulated genes of microglia during the acute SIV infection were used for the KEGG-ORA. **(D)** The hierarchical clustering of upregulated KEGG pathways for microglia in acute infection. **(E)** The gene-pathway networks between the specific pathways and human immunodeficiency virus 1. The complete pathways’ analyses with counts of mapped gene and p value were showed in Table S5. LOC114677644, LOC694372, LOC100426197, LOC719702 are the MHC class I molecules.

To better understand the changes in individual microglial clusters during acute SIV infection, we did a comparison between uninfected and acute infection samples for each microglial cluster **(Figure 3B).** We found that the genes that were related to MHC class I and interferon (IFN) signaling pathways were universally upregulated in microglial clusters during acute infection. It is well recognized that IFNs can block HIV/SIV replication,(^31–33^) and in this study we found that the key transcription factor in JAK-STAT pathway for IFN production, IRF9, was significantly upregulated in microglial cells. The increased fold-change of IRF9 ranged from 1.4 (Micro-11) to 8.3 (Micro-3), indicating this molecule was extensively upregulated in acute SIV infection. Furthermore, the genes encoding type I interferon inducible proteins (IFI genes), especially IFI6, IFI27 and IFI44 were also significantly upregulated in all microglial clusters. Those IFI genes are classified into interferon-stimulated genes (ISGs), which can be stimulated by the IFNs’ signaling to augment the restriction of HIV/SIV replication and cell entry. In addition, we also found that the IFNγ receptor (IFNGR1) was slightly increased in most microglial clusters (∼1.1-fold change) and highly increased in Micro-11 cluster (4.5-fold change). Other upregulated genes included those encoding for apolipoproteins (e.g., APOE, APOC1), lysosomal proteins (e.g., CTSC, CTSZ), complement components (e.g., C1QA, C1QB, C1QC), chemokines (e.g., CCL3, CCL4, CX3CR1), and proinflammatory cytokines (e.g., IL18, TNF, IL1B). Although most the aforementioned molecules were found to be upregulated in all microglial clusters, Micro-1, Micro-3, Micro-7, and Micro-11 were the clusters that had higher expression of those genes compared with Micro-0 and Micro-5 under acute SIV infection **(Figure S4A).** It should be noted that the Micro-5 cluster was found to have the highest expression of APOE and APOC in the uninfected condition as well as acute infection condition compared with other clusters **(Figure 3B).** APOE and APOC are also associated with neuroinflammation and neurodegeneration,(^34^) so their higher expression in Micro-5 indicated the activated nature of this cluster.

Surprisingly, different from the upregulated MHC class I molecules, the MHC class II molecules remained unchanged or downregulated for most microglial clusters. The downregulation of MHC class II molecules was more obvious in Micro-7 and Micro-11, in which the MHC class II molecules had higher expression than other clusters **(Figure 2A)**. The other downregulated genes were related to immunoregulation and microglial homeostasis (e.g., MERTK, TREM2, MRC1, FOLR2, SELPLG). This downregulation was significant in Micro-0, Micro-1, Micro-5, and Micro-7, but for Micro-3 and Micro-11 these genes’ expression was upregulated. When we compared the expression of these immunoregulatory or homeostatic molecules between different microglial clusters in acute SIV infection, we found that their upregulation in Micro-3 leads this cluster to have higher expression of those genes compared with other clusters **(Figure S4A)**. Intriguingly, CD4 gene expression was also downregulated in all microglial clusters (fold-change ranging from 1.1-1.3) and had highest expression in Micro-0 during the infection. CD4 serves as an important receptor for virus entry to the microglia(^35^) as well as the other targets of SIV/HIV infection, and its downregulation might indicate the defensive strategy of activated microglia in acute SIV infection. While CD4 downregulation by HIV and SIV infection of cells is known to occur through viral accessory proteins targeting the CD4 protein for degradation, (^36^) the downregulation of its transcript in all of the microglial clusters in acute infection likely is a result of cellular activation.

The genes that were significantly upregulated in microglial cells were then further associated with various pathways by using the Kyoto Encyclopedia of Genes and Genomes (KEGG) databases **(Figure 3C-3E).** All six clusters upregulated pathways related to antigen presentation and processing as well as cellular senescence. Micro-0 was found to upregulate the least number of pathological pathways compared with other microglial clusters **(Table S5)**, which suggests that the homeostatic status in Micro-0 was least altered by acute SIV infection. The other five microglial clusters were associated with more disease-related pathways under acute infection **(Figure S3)**. In general, these pathways could be categorized into five classes **(Figure 3D)**, including pathways related to antigen processing and presentation, viral infections, inflammation, cytosolic DNA sensing, and ribosomal activities. In the category of viral infections, we found the HIV-1 infection pathway, with sixteen enriched genes. Most genes upregulated in the HIV-1 infection pathway encoded MHC class I molecules, which connected the endocytosis, phagocytosis, and antigen presentation pathways to HIV-1 infection **(Figure 3E).**

The MEK2 protein kinase (MAP2K2) molecule, involved in mitogen-activated protein kinase (MAPK) pathway, also serves as the key molecule connecting cellular senescence and toll-like receptor (TLR) signaling with HIV-1 infection. MAP2K2 can trigger inflammation by phosphorylating the downstream kinases ERK1/2 (Extracellular Signal-Regulated Kinases 1 and 2) to translocate the transcription factor NF-κB and AP-1 to the nucleus for the expression of the genes encoding cytokines. The TLRs utilize those pathways to induce the production of proinflammatory cytokines. By comparing different TLRs, we found that TLR4, which can induce the expression of type I IFNs, was upregulated in all microglial clusters **(Figure 3C)** during acute SIV infection. TLR2 signaling, also acting through NF-κB and AP-1 to produce proinflammatory molecules (such as TNF-α, IL-1β, and IL-6), was upregulated in most microglial clusters. TLR7 which, as opposed to the cell-surface location of TLR4 and TLR2 is located on the endosomal compartment of the cells, and can specifically recognize single strand RNA (ssRNA) of HIV/SIV for the production of type I IFNs (^37–39^), was found to be upregulated particularly in Micro-3 (2.8-fold change). While the TLRs were extensively upregulated in microglial cells, Micro-7 and Micro-11 had the highest expression of TLR2, Micro-1 had the highest expression of TLR4, and Micro-3 and Micro-7 had the highest expression of TLR7 during acute SIV infection **(Figure S4A)**, indicating different microglial clusters might favor different TLR signaling pathways to produce proinflammatory cytokines. In summary, all of the upregulated pathways in acute SIV infection pointed to the pathways related to interferons and TLR-induced inflammatory cytokine production, highlighting the critical roles of them in microglia defense against acute SIV infection.

### The predominant CAM cluster phenotype in acute SIV infection was CD14^hi^CD16^low^

In the characterization of the predicted phenotypes for those two macrophage clusters, we found that Macro-6 had lower expression of the inflammatory molecules which were highly expressed in Macro-8 (e.g., APOE, CST3, MSR1). Instead, it had higher expression of the cell adhesion molecules (e.g., PECAM1 and integrins) **(Figure 4A and Table S2)**. In addition, Macro-6 was found to have high expression of CD16 (FCGR3) but low expression of CD14, and Macro-8 had high expression of CD14 but low expression of CD16 **(Figure 4B)**. In human blood, CD14^hi^CD16^low^ cells are described as inflammatory classical monocytes/ macrophages which are trafficked to sites of inflammation and/or infection, and CD14^low^CD16^hi^ cells are the patrolling non-classical monocytes/macrophages, which adhere and crawl along the luminal surface of endothelial cells.(^40^) In this study, we found that acute SIV infection shifted the dominant CAM phenotype from CD14^low^CD16^hi^ (Macro-6) to CD14^hi^CD16^low^ (Macro-8). Although Macro-8 appears to represent the classical inflammatory phenotype, it also had higher expression of the markers for anti-inflammatory M2-like macrophages (e.g., MRC1/CD206, CD163). Regarding differentiation, it was reported that the CD14^hi^CD16^low^ phenotype can be directly differentiated from precursor cells in bone marrow, and it is the obligatory precursor intermediate for CD14^low^CD16^hi^ phenotype in the blood.(^41^) Intriguingly, through trajectory analyses we also found that the CD14^low^CD16^hi^ CAMs (Macro-6) had a larger value of pseudotime compared to the CD14^hi^CD16^low^ CAMs (Macro-8) **(Figure 4C)**, suggesting CD14^hi^CD16^low^ cells might also be the precursor intermediate for CD14^low^CD16^hi^ cells in CNS.

**Figure 4.**
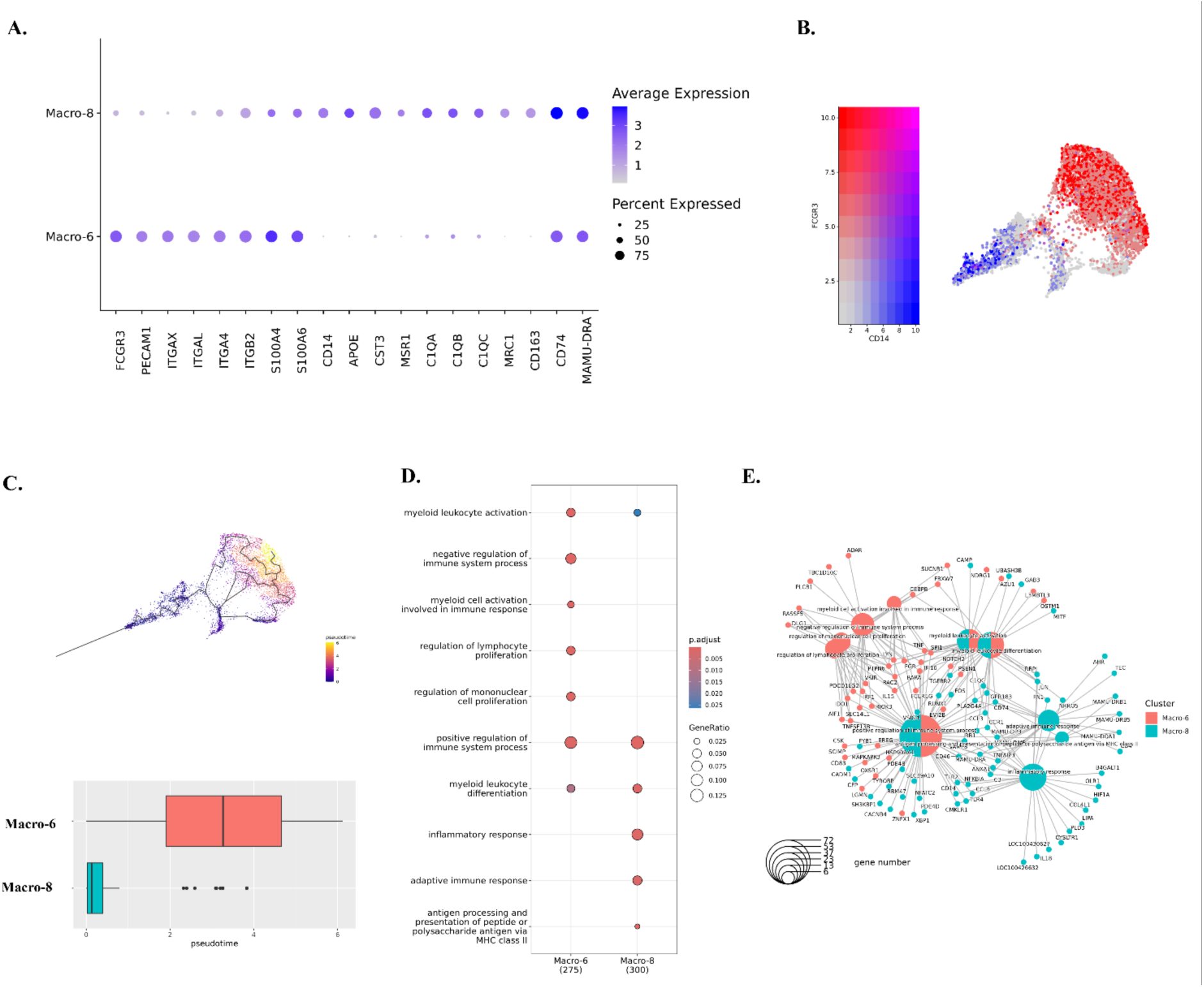
Characterizations of identified macrophage clusters. **(A)** Expression of DEGs in Macro-6 and Macro-8. The DEGs of each macrophage cluster were found by using Wilcoxon rank sum test for comparison. **(B)** The expression of CD14 and CD16 (FCGR3) in macrophage clusters. **(C)** The trajectory analyses for Macro-6 and Macro-8. The cell map indicated the trajectory paths, and the compassion of pseudotime between Macro-6 and Macro-8 was showed in boxplot. The larger value of pseudotime suggests the cells are more differentiated. **(D)** The characteristic biological processes in each macrophage cluster by Gene Ontology (GO) pathway analyses. **(E)** The networks between genes and biological pathways for macrophage clusters.

The GO results indicated that cells in both the Macro-6 and Macro-8 clusters participated in myeloid leukocyte activation and differentiation, which suggested these two phenotypes might originate or differentiate from infiltrating monocytes. Accordingto the opposite biological functions of CD14^hi^CD16^low^ and CD14^low^CD16^hi^ monocytes in the periphery, the Macro-6 cluster had both positive and negative regulations of the immune processes, however, the Macro-8 cluster lacked the negative regulation of the immune responses and thus had the potential to trigger inflammation **(Figure 4D)**. Furthermore, those two CAM clusters might also have the potential for interactions with infiltrating lymphocytes and adaptive immune responses, functions that were not found in the microglia clusters. For example, Macro-6 was predicted to regulate lymphocyte proliferation by secreting IL-15, which is an important stimulator for T/NK cells’ proliferation and activation. Macro-8 highly expressed MHC class II as well as other molecules, which can present the antigens for triggering adaptive immune responses **(Figure 4E)**. In summary, we identified both CD14^hi^CD16^low^ and CD14^low^CD16^hi^ macrophage phenotypes in the brains of rhesus macaque, and the CD14^hi^CD16^low^ cells (Macro-8 cluster) became the dominant phenotype in acute SIV infection, whereas CD14^low^CD16^hi^ cells (Macro-6 cluster) predominated in the uninfected condition.

### CAMs might be more activated in triggering immune responses under acute SIV infection than microglia

Consistent with the genes that were upregulated in all microglial clusters, most molecules related to MHC class I and IFN production were also significantly upregulated in both macrophage clusters **(Figure 5A and 5B)**. However, unlike microglial clusters, the IFI44L gene had extremely low expression in macrophage clusters. The changes of MHC class II molecules in macrophage clusters are different between Macro-6 and Macro-8. The expression of MHC class II molecules in Macro-6 remained unchanged except for the downregulation of MAMU-DPA, however, the expression of those genes in Macro-8 was extensively downregulated in acute SIV infection, which is consistent with the changes of activated microglial clusters. Intriguingly, we also found that the CAMs significantly upregulated the core genes for homeostatic microglia during the acute SIV infection **(Figure 5B)**. Given the myeloid lineage of microglia and CAMs, the similar behaviors in response to acute SIV infection might be expected (e.g., APOE and SPP1), but it was surprisingly that the homeostatic genes for microglia (e.g., P2RY12, GPR34) and other genes characterizing microglia (MERTK, SELPLG) that were downregulated or remained unchanged in microglial clusters during acute SIV infection were increased in CAMs, especially Macro-6. We found that that P2RY12 expression was upregulated 4-fold, GPR34 4.5-fold, SELPLG 2.4-fold, and MERTK 3.7-fold in the Macro-6 cluster. Although Macro-6 highly upregulated those molecules, the expressions of microglial homeostatic core genes were still higher in Macro-8 during acute infection **(Figure S4B)**. Furthermore, while Macro-6 was characterized as CD14^low^CD16^hi^ macrophages, and Macro-8 was characterized as classic CD14^hi^CD16^low^ macrophages, once they were in acute infection condition, the CD14^hi^CD16^low^ cells upregulated CD16 (1.5-fold), whereas the CD14^low^CD16^hi^ cells upregulated CD14 (3.3-fold). This again highlights the plasticity of myeloid cells.

**Figure 5.**
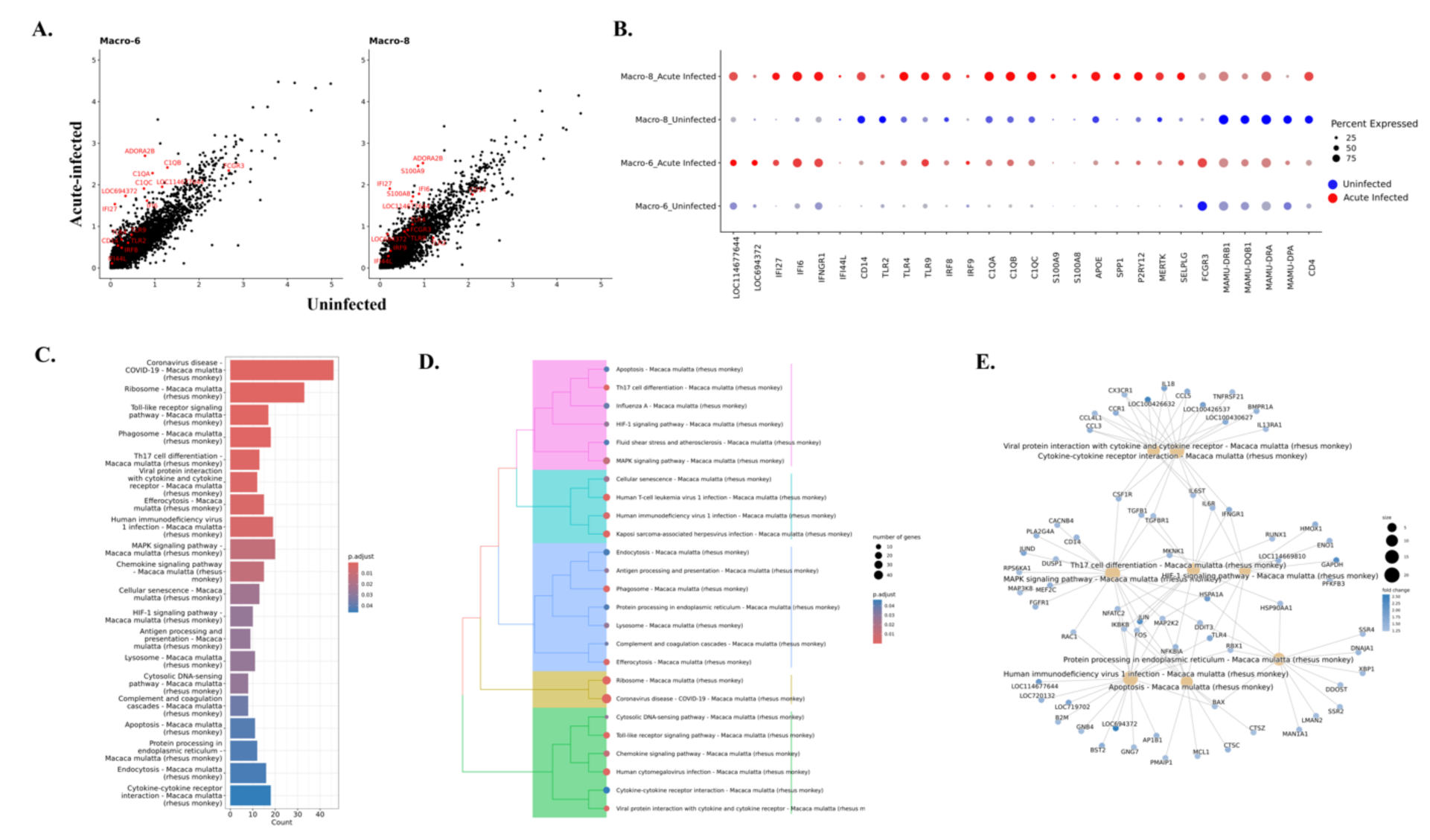
Altered gene expressions and pathways in macrophage during acute SIV infection. **(A)** The comparison of the gene expression levels between the acute-infected animals and uninfected animals for macrophage clusters. Some of the significantly changed genes were labeled in red. **(B)** Selected DEGs in uninfected animals (blue dots) and animals with acute infection (red dots) for each macrophage cluster. **(C)** Top 20 upregulated pathways in macrophages from acute infection animals. Overall upregulated genes with adjusted p values of less than 0.05 in macrophage were used for the Kyoto Encyclopedia of Genes and Genomes (KEGG) pathways analyses. **(D)** The hierarchical clustering of upregulated KEGG pathways of macrophage in acute infection. **(E)** The gene-pathway networks between the specific pathways and human immunodeficiency virus 1 pathway. LOC114677644 and LOC694372 are the MHC class I molecules.

The elevation of CD14 in Macro-6 was also accompanied by the increased expression of TLR4, which is the coreceptor for CD14 in inducing pro-inflammatory signaling. In addition, more genes related to the inflammatory pathways (e.g., TLR2, TLR9, IRF8, IRF9, C1QA, C1QB, C1QC) were found to be increased in both macrophage clusters **(Figure 5B)**. The fold-change of most inflammatory molecules was higher in Macro-6 (e.g., C1QA, C1QB, and C1QC had a 2-fold change in Macro-6 but only ∼1.2-fold change in Macro-8) but the overall expression level was still higher in Macro-8 compared to Macro-6 **(Figure S4B)**. Macro-8 also significantly upregulated more molecules related to innate defense, such as S100A8 (fold-change: 2.3) and S100A9 (fold-change: 2.9), which was not observed in Macro-6. In summary, these results suggest that the immunosuppressed phenotype of Macro-6 might differentiate toward inflammatory phenotype, and the pre-activated phenotype of Macro-8 upregulated more inflammatory molecules and signaling pathways during the acute infection.

The KEGG gene enrichment analyses for macrophages showed more upregulated inflammatory pathways compared to microglia **(Figure S5 and Table S5)**. Overall, the macrophages not only enhanced the pathways that were found to be upregulated in microglia, but also augmented other inflammation-related pathways, for example, the interaction between viral proteins and cytokine/chemokine receptors and Th17 cell differentiation **(Figure 5C and 5D)**. In the characterization of the CAMs by their featured pathways, we found that these macrophages might have more interactions with the T cells than microglial cells. When we further examined the genes that were upregulated during acute SIV infection in the KEGG pathways, we found that they were enriched in the ability to induce Th17 differentiation during acute SIV infection. The genes that are mapped to this pathway included IL-6 and TGFB **(Figure 5E)**, which were reported as key cytokines secreted by macrophages to trigger differentiation of Th17 cells.(^42, 43^) Given the pleiotropic functions of those two cytokines, their increase might not be necessarily correlated with Th17 differentiation, but their increased expression in perivascular macrophages responding to acute SIV infection has been confirmed in rhesus macaque.(^44^) Even though more inflammatory or immunological pathways were upregulated in macrophages compared to microglia, the key pathway that connects them with HIV-1 infection might also involve the NF-κB signaling. In the gene-pathway connections **(Figure 5E)**, multiple molecules and kinases for NF-κB signaling (e.g., IKBKB, JUN, FOS, MAP2K2, NFKBIA, TLR4) connected the HIV-1 infection and other inflammatory or anti-viral pathways.

### The genes that were upregulated in Micro-3 and Macro-8 clusters in response to acute SIV infection are associated with multiple neurological diseases

The microglial and macrophage clusters that were widely activated, with increased expression of proinflammatory cytokines and chemokines in acute SIV infection, might induce toxicities not only for the virus but also for neurons and other cells in the CNS. Here, we found that one microglial cluster and one macrophage cluster may be strongly related to neurological disorders such as HAND. The upregulated genes analyzed with KEGG gene enrichment analyses for each cluster revealed that there was a category of pathways including those leading to multiple neurocognitive disorders in both the Micro-3 and Macro-8 clusters **(Figure 6A**). Prion disease, Parkinson disease, Alzheimer disease, Huntington disease, and amyotrophic lateral sclerosis were in the list of this category. In Micro-3, all of those five neurodegeneration-associated pathways have extremely low adjusted p-value (<0.002); in Macro-8 the Huntington disease and amyotrophic lateral sclerosis had a slightly higher adjust p-value (∼0.006). The mapped genes in those pathways were related to mitochondrial functions, and we found that the fold-change of the associated genes in Macro-8 was higher than the Micro-3 **(Figure 6B)**.

**Figure 6.**
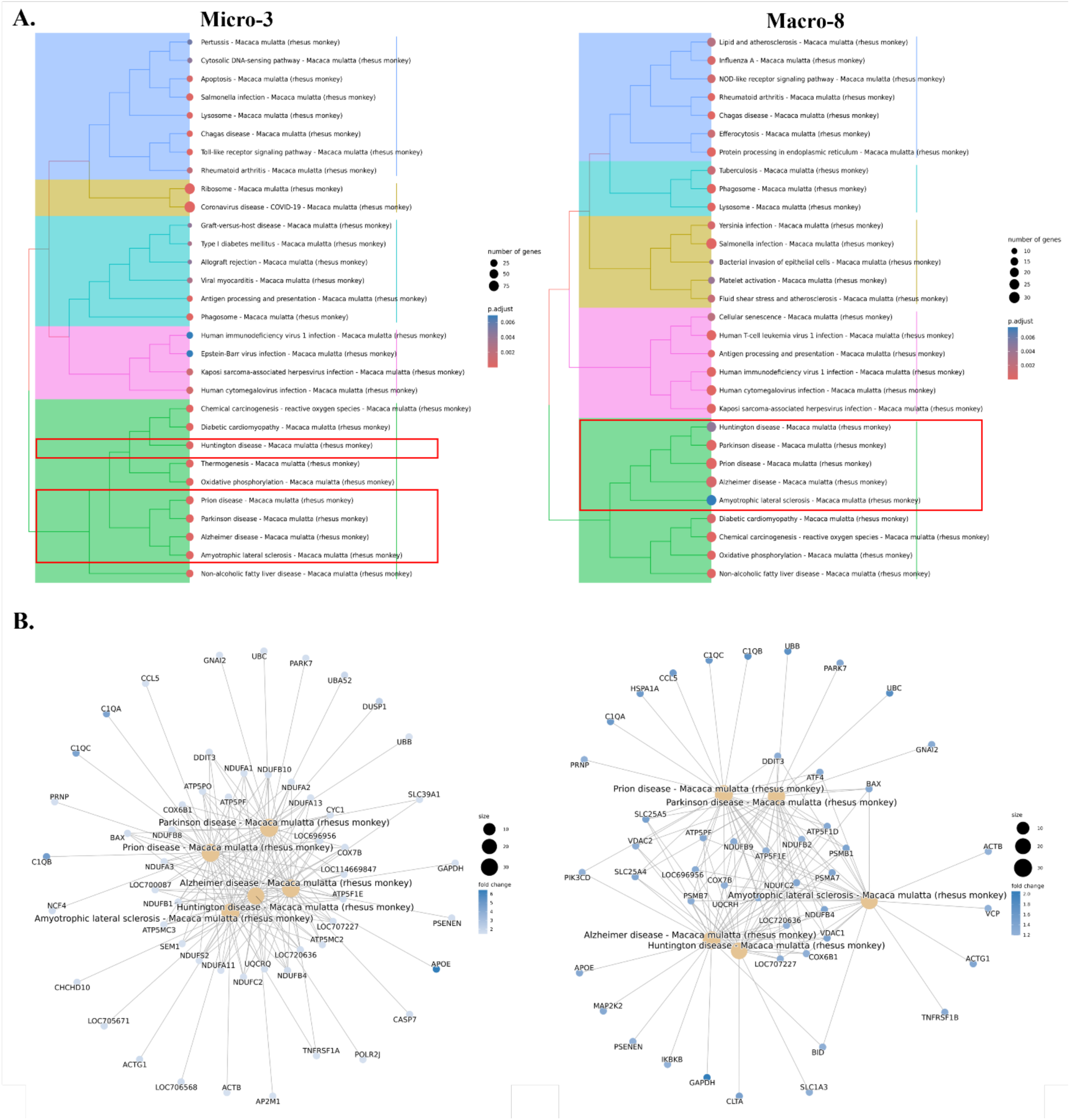
Potential neurodegenerative pathways associated with Micro-3 and Macro-8. **(A)** In the hierarchical clustering of the upregulated KEGG pathways for each cluster during the acute SIV infection, there was a category for Micro-3 (left) and Macro-8 (right) which enriched with multiple neurodegenerative diseases. There were five neurological diseases identified in both clusters, including Prion disease, Parkinson disease, Alzheimer disease, Huntington disease, and Amyotrophic lateral sclerosis. **(B)** The genes that were associated with each mapped neurodegenerative pathway for Micro-3 (left) and Macro-8 (right).

To further confirm the specificity of neurodegenerative molecules in Micro-3 and Macro-8 during the acute SIV infection, we then compared the expression changes for a number of these genes in all microglia and macrophages under uninfected conditions and infected conditions. Consistent with the above, we found that those genes were significantly upregulated in Micro-3 and Macro-8 in acute SIV infection **(Figure 7A and Figure 7C)**. However, the expression of these genes barely changed when we compared them in all microglial cells **(Figure 7B)**. Although genes that are related to the mitochondria functions seemed to be upregulated when all macrophages in infected and uninfected conditions were compared **(Figure 7D)**, the overall changing fold was not as prominent as in the Macro-8 cluster. All of these results suggested that the Micro-3 and Macro-8 clusters might be strongly associated with the induction of neurological disturbances when activated in acute SIV infection.

**Figure 7.**
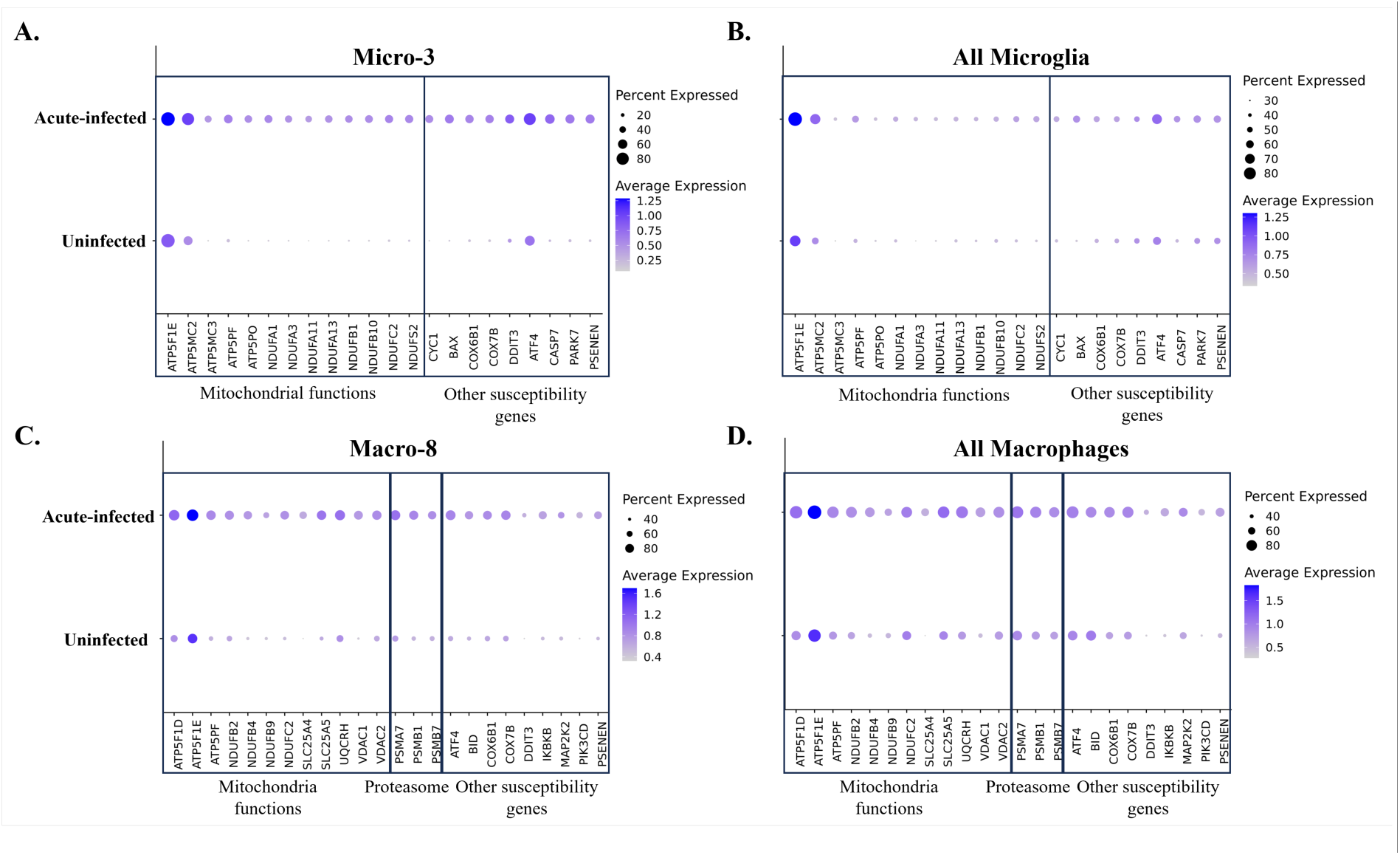
Comparison of the susceptibility genes for neurodegenerative disorders in uninfected animals and acute-infected animals. The comparison was conducted for the cells in **(A)** Micro-3 cluster, **(B)** all microglial clusters, **(C)** Macro-8 cluster, **(D)** and all macrophage clusters separately. The susceptibility genes mapped to the neurodegenerative pathways were compared, which includes the genes related to the mitochondria and proteasome functions.

### Apoptotic resistance was not initiated during acute SIV infection in myeloid cells in the brain

HIV was reported to upregulate the expression anti-apoptotic molecules and downregulate pro-apoptotic molecules to enable the survival of infected host cells.(^25, 26, 45, 46^) Those anti-apoptotic molecules (e.g., BCL2, BFL1, BCL-XL, MCL-1) and pro-apoptotic molecules (e.g., BAX, BIM, BAK1, BAD) are typically found in BCL-2 family. While microglia and macrophages are typically thought to be long-lived, the mechanism of survival following HIV infection has not been extensively studied. To examine if HIV infection of myeloid cells in the brain could lead to the establishment of the virus reservoir by blocking apoptosis, we first compared the expression of anti-apoptotic and pro-apoptotic molecules between macrophages and microglia in acute-infected animals and uninfected animals **(Figure 8A)**. However, we did not observe remarkable changes in the RNAs encoded by these molecules in macrophages and microglia during acute infection. We did observe the anti-apoptotic molecule BCL2 was downregulated, and the pro-apoptotic molecule BAX was upregulated, suggesting that the macrophages and microglia are prone to undergo apoptosis *in vivo* upon SIV infection. Then we further investigated the change of these molecules in individual myeloid cell cluster **(Figure 8B)**. The BCL2 gene was downregulated in most clusters, and the cluster which did not downregulate BCL2 (Macro-6) was found to maintain its expression without change. On the other hand, the pro-apoptotic gene, BAX was widely upregulated in most cell clusters. For other anti-apoptotic gene expressions, we found that MCL1 was upregulated, except in homeostatic Micro-0 and Micro-1, CFLIP (CFLAR) was upregulated in Micro-3 and Micro-11, and others remained unchanged or slightly downregulated in infection. The pro-apoptotic molecules were generally upregulated, for example, BAK1 was upregulated in Micro-3, Micro-7, and Micro-11, BID was upregulated in Micro-3 and Macro-8, and PMAIP1 was upregulated in Micro-11. However, a known pro-apoptotic molecule for microglia and macrophage, BIM (BCL2L11), was found to be downregulated in Macro-6 and Macro-8. Interestingly, we found that another anti-apoptotic molecule CD5L, which is not included in BCL-2 family, was highly upregulated in Micro-11.

**Figure 8.**
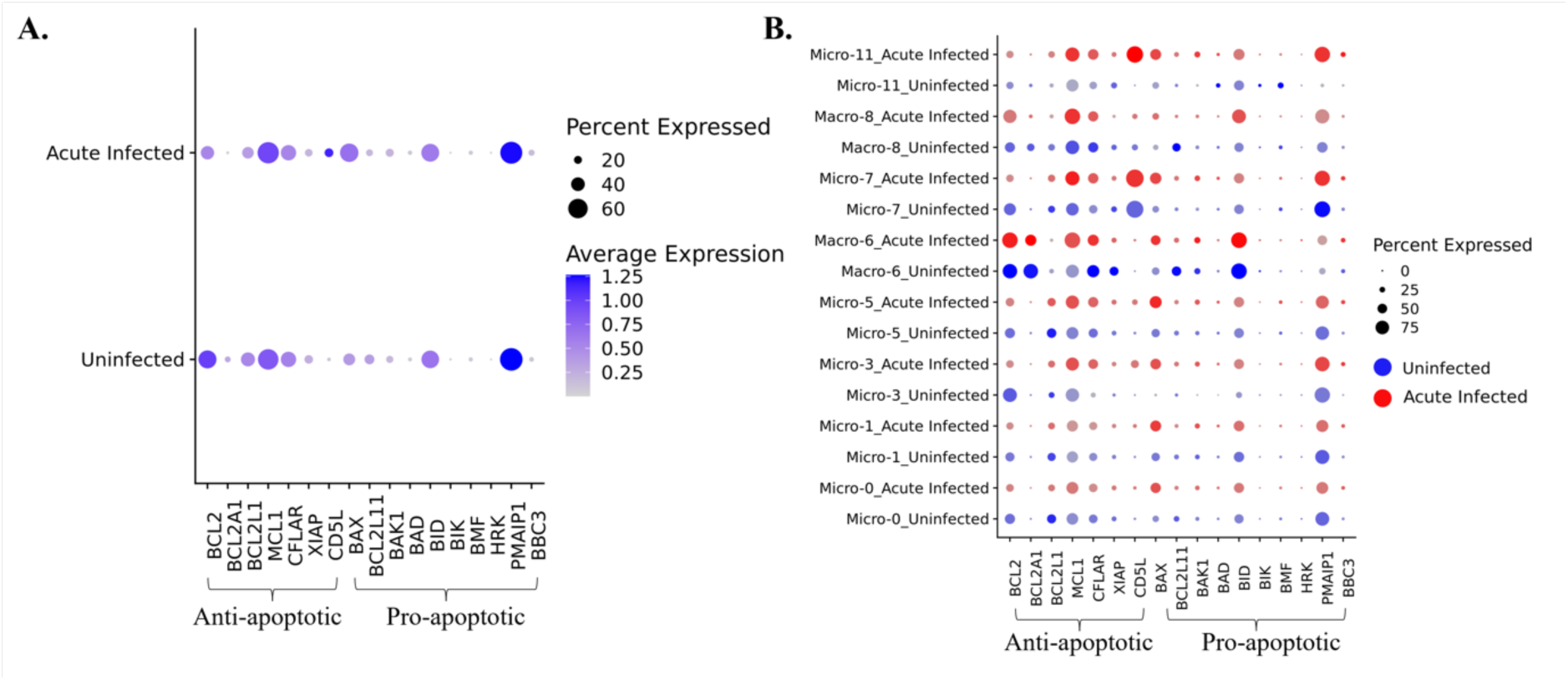
Changes of anti-apoptotic and pro-apoptotic genes in brain myeloid cells due to acute SIV infection. **(A)** The average expression of anti-apoptotic and pro-apoptotic genes in acute infection or uninfected animals was compared in all macrophages and microglia combined. **(B)** The average expression of anti-apoptotic and pro-apoptotic genes in acute infection (red dots) or uninfected (blue dots) animals was compared in each macrophage and microglial cluster.

## DISCUSSION

Microglia are the major innate immune cells in the brain and serve multiple functions. They are active in protecting the brain from invading pathogens, clearing damaged synapses as well as dying cells, and promoting neuron development. While not as extensively characterized as neurons, they are indeed heterogeneous, and with the advent of scRNA-seq our studies and those of others have identified a number of classes of microglia.(^47–50^) To maintain a steady state while gaining some immunogenic properties, microglia express several homeostatic genes. Downregulation and/or mutations of these homeostatic genes can lead to uncontrolled neuroinflammation, neurotoxicity and multiple neurocognitive diseases.(^51–55^) In this study, we found multiple populations of microglia which changed their proportions during acute SIV infection. During acute SIV infection overall activation was observed, with a lower expression of homeostatic genes and a decreased proportion of cells with a homeostatic phenotype. In addition, we found the major homeostatic microglial cluster, Micro-0, was enriched in the TGFβ signaling pathway, which was not observed for the other microglial clusters. Based on others’ findings, the silencing of TGFβ signaling in microglia resulted in loss of microglial ramification and the upregulation of inflammatory markers without any external stimulus relative to wild-type microglia.(^30, 56^) Considering the pleiotropic functions of TGFβ signaling in microglial cells,(^57–59^) the more active TGFβ pathway in Micro-0 suggested this homeostatic microglial cluster might be more active in stimulating microglial differentiation and regulating the activation of other microglial clusters.

Although the proportion of cells in Micro-0 remarkably decreased during acute SIV infection, this was still the dominant microglial population (consisting of 44.1% of microglial cells in acute infection, compared to 64.1% in uninfected animals) in the brain, suggesting that microglial activation was still under some control during the acute infection stage. In addition, Micro-0 did not show prominent signs of activation during this stage, which could be attributed to their high activity in TGFβ signaling. The Micro-1 cluster, with high expression of microglial core genes but lack of TGFβ signaling pathway enrichment, was much more inducible in terms of gene expression during acute SIV infection than the homeostatic Micro-0 cluster. More pathogenic and inflammatory pathways were upregulated for Micro-1 during acute SIV infection **(Figure S3)**, highlighting the importance of TGFβ signaling pathway in regulating microglial activation.

Micro-1, similar to Micro-0, also decreased its proportion in acute infection, comprising 27.1% of the myeloid cells versus 32% in the uninfected animals. These decreases were accompanied by prominent changes in the proportions of other microglial populations in acute infection, as Micro-3, −5, −7, and −11 all increased their representation. For CAMs the dominant cluster changed from Macro-6 (65% in uninfected animals) to Macro-8 (72.9% in acute SIV infection). All these were characterized by increased signs of cellular activation.

In uninfected brains, the activated microglial clusters constituted a low but noticeable presence (<5% proportion). Those activated microglia cells have been reported to exist in the healthy brain at different anatomic locations and are involved in diverse neurological events compared with homeostatic microglia.(^60–62^) From scRNA-seq experiments, it is difficult to determine the biological meaning of such low proportion cell clusters. Therefore, to better interpret the changes in gene expression during the acute SIV infection, we did the comparisons in multiple ways.

We found that MHC class I genes and genes related to interferon production were significantly upregulated in both microglial and macrophage clusters during acute SIV infection. However, the expression levels were different between individual clusters, suggesting discrepancies of responses for each cluster. For virus infection, MHC class I molecules are key to the host defense for their ability to present virus proteins in the infected cells to the cytotoxic T/NK cells in MHC class I manner. (^63^) The significant upregulation of MHC class I molecules in all myeloid cell clusters also indicated that the reaction to initial SIV infection might not be limited to the low proportion of microglia and macrophages infected with SIV, and that the bystander cells also showed reactivity to the infection. This was further supported by the transcriptional changes in these bystander cells (**Table S3**).

The expression of type I interferons, especially IFNβ(^64, 65^) has been found to be widely upregulated in acute SIV infection. In response to the production of IFNβ, the downstream ISGs that are induced by type I IFNs to amplify the antiviral effects have been found to be upregulated in blood and lymph nodes during the acute SIV/HIV infection in rhesus macaque and human.(^66, 67^) Our recent and prior studies of microglial responses to acute and chronic SIV infection in the brain(^50, 68^) also highlighted that numerous ISGs were significantly upregulated in the myeloid cells of the brain and might contribute to the HIV/SIV associated brain damages. Therefore, the high expression of ISGs starting at an early stage of SIV infection can extend to chronic infection in the brain’s myeloid cell populations.

We note that one of the important pathways for type I IFN production is TLR signaling, and we found that TLR2 and TLR4 were widely upregulated in microglial and CAM clusters. TLR7, which can specifically bind with ssRNA from SIV, was upregulated in Micro-3, Micro-5, and Micro-11, and TLR9 which can bind with the CpG motif in DNA was widely upregulated in CAM clusters. All of these TLRs were reported to have the ability to recognize and bind with HIV,(^69–71^) although TLR7 might be the primary target.(^37^) The presence of TLR4, TLR7, and TLR9 in endosomal compartments endows them the ability to recognize viral nucleic acids in the cells, and indeed their signaling pathways are related to the production of IFNs.(^72–74^) TLR signaling can trigger the transcription of numerous inflammatory cytokines (e.g., TNF-α, IL-6, IL-1β, etc), which were also found upregulated in microglial or macrophage clusters under acute SIV infection. All of those cytokines can serve as initiator or the products for the pathways associated with NF-κB and AP-1 signaling, which also has been found to be modulated by HIV/SIV for their establishment and reactivation from latency.(^75–79^)

CNS-associated macrophages, found in the interface between the parenchyma and the circulation, represent an important myeloid cell population in the brain distinct from microglia. Previously, all CAMs were thought to be derived from the monocytes in the circulation, but recent findings showed that CAMs are highly heterogenous. Some phenotypes might originate from the yolk sac and can be self-replenishable like microglia, which are different from the CAMs differentiating from the circulating monocytes.(^80, 81^) From the transcriptional perspective, the yolk sac-derived CAMs also share more similarities with microglia, and they are hard to separate based on their transcriptomic profiles.(^80^) Given the distance between CAM clusters and the microglial clusters on the UMAP and the low expression of microglial core genes in CAM clusters, Macro-6 and Macro-8 were more likely to be derived from the circulating monocytes. Like the two main phenotypes in the periphery, Macro-6 had a CD14^low^CD16^hi^ phenotype with immunosuppressive properties and Macro-8 had a CD14^hi^CD16^low^ phenotype with proinflammatory properties. During acute SIV infection, Macro-6 upregulated more inflammatory molecules which were already highly expressed in Macro-8, indicating the Macro-6 could be activated and polarized toward the inflammatory CD14^hi^CD16^low^ phenotype. This observation is consistent with a report about the transcriptomic convergency of peripheral CD14^++^CD16^+^ and CD14^+^CD16^++^ population under the SIV infection.(^82^)

Intriguingly, macrophages also increased the expression of microglial core genes during the acute SIV infection. In a different system, it was found that the monocyte-derived macrophages in the retina adopted microglia-like gene expression during the degeneration processes,(^83^) which was similar to what we found for the CAMs in acute SIV infection. However, the reasons why CAMs share more similarities with microglia in response to stress or neurodegenerative diseases are still unknown. Although the CAM clusters synchronized the changes for most immunoreactive molecules, other immunoreactive molecules might change differently. For example, most MHC class II molecules that were not changed or slightly upregulated in Macro-6 were downregulated in Macro-8. The repression of MHC class II molecules in professional antigen-presenting cells (APCs) was also observed during HIV infection in humans, and serves as one of the immunodeficiency mechanisms of CD4+ T cells in AIDS.(^84–87^) Therefore, the unchanged MHC class II in Macro-6 suggested that this cluster might be less likely to be affected by SIV-induced immunodeficiency at least in acute infection. They may still maintain the ability to trigger the activation and differentiation of CD4+ T cells for host defense against SIV virus.

The activation of microglia and CAMs in the brain is essential for protecting against viral infection, however, the pro-inflammatory cytokines and cytotoxic molecules secreted by them might also lead to the damages for neurons as well as other supportive apparatus in CNS. Therefore, overreactive microglia and especially monocyte-derived CAMs were thought to play a central role in HAND before the era of efficacious antiretroviral therapy.(^24, 88^) By enriching the upregulated genes in each microglial or macrophage cluster into the various pathways, we found that Micro-3 and Macro-8 clusters were linked with numerous neurological diseases, including Alzheimer’s, Huntington’s, and Parkinson’s diseases during acute SIV infection. These changes CAMs and microglia also have potential in the pathogenesis of HAND by various disease mechanisms associated with neurodegeneration.(^24^) When we further identified the upregulated genes that were linked with those neurological disorders, we found that, the genes related to mitochondrial respiration were significantly upregulated in both Micro-3 and Macro-8. *In vitro*, HIV infection of macrophages has been shown to alter their mitochondrial energetic profiles, with the specific changed dependent on the stage of infection.(^89^)

Given the unmet need for the validated biomarkers of HAND in clinical settings,(^90^) some studies have sought to identify various proinflammatory cytokines and molecules associated with protein misfolding as biomarkers for HAND,(^91, 92^) and other studies explored the specific myeloid cell phenotypes for serving as biomarkers.(^88, 93^) Both HIV itself and the antiretroviral drugs used to treat HIV can affect mitochondria, and mitochondrial pathways are key suspects in the resulting neuropathogenesis of HIV and HAND.(^94^)

Although only Micro-3 and Macro-8 showed enrichment in multiple neurological disorder pathways, all the myeloid cell clusters were associated with the cellular senescence pathway, which might be another potential factor triggering or dampening the neurological disturbances.(^95, 96^) The potential effect of cell aging caused by acute SIV infection might not only happen in microglia or macrophages but extend to neurons as well as other brain cells. Indeed, while very few myeloid cells were directly infected, the bystander effects of infection were widely manifested. Furthermore, the inflammatory microglia and CAMs resulting from virus infection are also likely to trigger or speed up cerebral aging.(^97^) Indeed epigenetic studies have found that HIV infection leads to an advancement of predicted biological age in blood cells as well as in the brain, and is associated with reduction in brain gray matter and cortical thickness.(^98–101^)

Both HAND and the establishment of HIV/SIV reservoir in CNS can occur irrespective of antiretroviral therapy.(^102, 103^) Macrophages and microglia cells are believed to serve as major HIV reservoir in the CNS given that they are prime targets for the virus and their known longevity.(^14^) Many pathways could be altered by HIV infection allowing for the long-term survival of the infected macrophages and microglia, and the anti-apoptotic or pro-apoptotic molecules in BCL-2 families were found to regulate the survival of macrophages and/or microglia in HIV infection.(^25, 26, 104, 105^) However, we did not observe a wide and significant upregulation of anti-apoptotic molecules or downregulation of pro-apoptotic molecules in myeloid cell populations identified in this study. The cluster that upregulated anti-apoptotic genes also upregulated pro-apoptotic genes, so it is difficult to identify a specific cluster that might facilitate the establishment of a SIV, and by analogy HIV, reservoir. From the average expression of the anti-apoptotic and pro-apoptotic molecules in BCL-2 families, the overall trend for the myeloid cells of the brain under acute SIV appears to be the enhancement of pro-apoptosis and suppression of anti-apoptosis, and the means in which reservoir cells survive long-term in unknown.

Still there might be other molecules in myeloid cells that can also regulate the apoptotic pathways. For example, we found upregulation of CD5L, which is also known as apoptosis inhibitor of macrophage (AIM). CD5L can support the survival of macrophages when the cells are challenged with infections or other dangers.(^106^) Although anti-apoptosis(^107^) is the most well-recognized function for this molecule, little is known about the intracellular mechanisms underlying CD5L regulation of apoptosis. Whether this molecule could be modulated during the SIV/HIV for modulating the virus reservoir in macrophages or microglia needs further investigations.

Limitations to our study include the number of animals examined, study of a single time point during the acute infection period, the lower sensitivity of scRNA-seq, the lack of spatial assessment within the brain of where these populations of cells are present, and potential differences between SIV infection of rhesus monkeys and HIV infection of people. However, studies of the brain in people are largely limited to post-mortem studies and lack control over conditions. Yet many considerations limit the number of nonhuman primates used for terminal experimental studies. The deposition of sequence data and metadata in publicly accessible databases from our studies and others’ enables the building of larger analyses with more subjects. These data can be useful in meta-analyses across models and disease states.(^108^) The ability to purify microglia and macrophages from the brains and the ability to examine thousands of cells from each animal is an advantage enabling the identification and study of less prevalent populations. The continued development of spatial transcriptomics, sequencing methods and other technological and analytic advances will also help alleviate these limitations.

In conclusion, by performing scRNA-seq to assess the brain myeloid cells in rhesus macaques, we identified six microglial clusters and two macrophage clusters. In response to the acute SIV infection of a small proportion of cells, all myeloid clusters upregulated the genes related to MHC class I molecules and IFN signaling, which also served as the key connections for other cellular responses to the HIV/SIV infection. The activated microglial and macrophage clusters with more upregulated inflammatory cytokines increased their proportions and the homeostatic or immunosuppressive myeloid clusters decreased their proportions during acute SIV infection. Among the activated clusters, both a microglial cluster and a macrophage cluster were identified that exhibited dysregulation of genes associated with pathways linked to neurodegenerative disorders. Changes in all of the microglial clusters may contribute to worsening neurological health due to their enhanced ability to produce inflammatory molecules and involvement in cellular senescence.

## MATERIALS AND METHODS

### Animals

The six male adult rhesus macaques used in this study were purchased from PrimGen (Hines, IL) and New Iberia (LA), and tested negative for the indicated viral pathogens: SIV, SRV, STLV1, Herpes B-virus, and measles; and bacterial pathogens: salmonella, shigella, campylobacter, yersinia, and vibrio. Macaques were housed in compliance with the Animal Welfare Act and the Guide for the Care and Use of Laboratory Animals in the NHP facilities of the Department of Comparative Medicine, University of Nebraska Medical Center (UNMC). The primate facility at UNMC has been accredited by the American Association for Accreditation of Laboratory Animal Care international. The UNMC Institutional Animal Care and Use Committee (IACUC) reviewed and approved this study under protocols 19-145-12-FC and 16-073-07-FC. Animals were maintained in a temperature-controlled (23 ± 2° C) indoor climate with a 12-h light/dark cycle. They were fed Teklad Global 25% protein primate diet (Envigo, Madison, WI) supplemented with fresh fruit, vegetables, and water *ad libitum*. The monkeys were observed twice daily for health status by the animal care and veterinary personnel. Three of the six animals were intravenously inoculated with a stock of SIV_mac_251 to establish acute SIV infection (93T, 94T, and 95T). The other three macaques were uninfected (92T, 104T, and 111T) and used as control. Virus stocks were provided by the Virus Characterization, Isolation and Production Core at Tulane National Primate Research Center.

### Viral loads

To determine the viral load in plasma, the blood of infected animals (93T, 94T and 95T) was collected at 7-day and 12-day post-inoculation of SIV. The EDTA-anticoagulated plasma was separated from blood by centrifugation. Specimens of brain and lymphoid organs were taken for determination of viral load in tissues. Plasma SIV RNA levels were determined using a gag-targeted quantitative real time/digital RT-PCR format assay, essentially as previously described, with six replicate reactions analyzed per extracted sample for assay threshold of 15 SIV RNA copies/ml.(^109^) Quantitative assessment of SIV DNA and RNA in tissues was performed using gag-targeted nested quantitative hybrid real-time/digital RT-PCR and PCR assays, as previously described.(^109, 110^) SIV RNA or DNA copy numbers were normalized based on quantitation of a single copy rhesus genomic DNA sequence from the CCR5 locus from the same specimen to allow normalization of SIV RNA or DNA copy numbers per 10^6^ diploid genome cell equivalents, as described.(^111^) Ten replicate reactions were performed with aliquots of extracted DNA or RNA from each sample, with two additional spiked internal control reactions performed with each sample to assess potential reaction inhibition. The viral load in plasma, lymphatic tissues and brain are shown in **Figure S1** and **Table S1**.

### Isolation of myeloid cells in the brain

Twelve days after viral inoculation, necropsy was performed on deeply anesthetized (ketamine plus xylazine) animals, following intracardial perfusion with sterile PBS containing 1 U/ml heparin. Brains were harvested and approximately half of the brain was then taken for microglia/macrophage isolation. Microglia/macrophage-enriched cellular isolation was performed using our previously described procedure.(^112^) Briefly, the brain was minced and homogenized in cold Hank’s Balanced Salt Solution (HBSS, Invitrogen, Carlsbad, CA). After being centrifuged, the brain tissue was then digested at 37° C in HBSS containing 28 U/ml DNase I and 8 U/ml papain for 30 minutes. After digestion, the enzymes were inactivated by addition of 2.5% FBS, and the cells centrifuged and resuspended in cold HBSS. The cell suspension was mixed with 90% Percoll (GE HealthCare, Pittsburg, PA) and centrifuged at 4° C for 15 minutes at 550 x g. The microglia/macrophage pallet at the bottom was resuspended in HBSS and passed through a 40 μm strainer to remove cell clumps and/or aggregates. Cells were again pelleted by centrifugation and resuspended in RBC lysis buffer for 3 minutes to eliminate any contaminating red blood cells. A final wash was performed before the resulting cells were quantified on both a hemocytometer and Coulter Counter Z1. The isolated cells were then sorted for scRNA-seq. The cells in samples 92T, 93T, 94T, 95T were sorted and sequenced right after the isolation, but the cells in samples 104T and 111T were cryopreserved before the cell sorting and scRNA-seq. The methods for cryopreservation followed our previous study, which was found to maintain the vast majority of the transcriptomic features of fresh isolated microglia/macrophages.(^112^) Specifically, after isolation as above, the cells were centrifuged at 4° C at 550 x g for 5 minutes and the supernatant was removed. The pellet was dissociated by tapping and then resuspended by the dropwise addition of a solution of 4° C 10% DMSO in FBS at a concentration of 10^6^ cells per milliliter. Cells were transferred to cryopreservation tubes and slowly controlled freezing at −80° C. After 24 hours, cryotubes were transferred to liquid nitrogen for long-term storage.

### Single cell preparation and RNA sequencing

For fresh brain isolates, cells were washed in PBS, and first stained with UV-blue live/dead. Cells we then washed, resuspended in MACS buffer with 0.1% BSA, and counted. The cells were then stained with non-human primate CD11b microbeads and CD45 microbeads (Miltenyi, San Diego, CA, USA). Four hundred million cells were reconstituted in 320 μL of MACS buffer and reacted with 180 μL of CD11b and 40 μL of CD45 microbeads at 4° C for 15 minutes. After incubation, cells were washed, resuspended in 1 ml of MACS buffer, and loaded on MACS Separator LS columns. The double enriched fractions were collected and counted, and then stained with antibody cocktails including BV711-labeled anti-CD20 antibody, BV421-labeled anti-CD3 antibody, BV605-labeled anti-CD11b antibody and PE-labeled anti-CD45 antibody (Biolegend, San Diego, CA) for 45 minutes at 4° C. Cells were washed with e-bioscience flow cytometry staining buffer and sorted on Aria2 flow cytometer (BD Biosciences, San Jose, CA, USA). The selection of cells was based on the size, singlet, live and the expression of CD20, CD3, CD45, and CD11b. The CD20 positive cells were all excluded, and the CD20 negative cells that were positive for either CD45 or CD11b or both CD45 and CD11b were collected for scRNA-seq library preparations.

Samples of cryopreserved cell isolates, stored in liquid nitrogen described above, were rapidly thawed in a 37° C water-bath. The cell recovery procedures were well described in our previous publications.(^112^) After the recovery, cells were washed and counted by Coulter Counter Z1. Once cell concentration was known, cells were transferred to ice-cold PBS, and followed the aforementioned procedures for CD45 and CD11b enrichment and FACS sorting.

Post-sorting, isolates were concentrated to approximately 1000 cells per µL, and assessed by trypan blue for viability and concentration. Based on 10× Genomics parameters targeting 8000 cells, the ideal volume of cells was loaded onto the 10× Genomics (Pleasanton, CA, USA) Chromium GEM Chip and placed into the Chromium Controller for cell capturing and library preparation. This occurs through microfluidics and combining with Single Cell 3’ Gel Beads containing unique barcoded primers with a unique molecular identifier (UMI), followed by lysis of cells and barcoded reverse transcription of RNA, amplification of barcoded cDNA, fragmentation of cDNA to 200 bp, 5’ adapter attachment, and sample indexing as the manufacturer instructed with version 3 reagent kits. The prepared libraries were then sequenced using Illumina (San Diego, CA, USA) Nextseq550 and Novaseq6000 sequencers. The sequences have been deposited in NCBI GEO (accession number GSE253835).

### Bioinformatics

Sequenced samples were processed using the 10× Genomics Cell Ranger pipelines (7.1.0). Specifically, the scRNA data was demultiplexed and aligned to the customized Mmul10 rhesus macaque reference genome (NCBI RefSeq assembly) which was combined with a chromosome representing the SIV genome. For this, overlapping PCR products derived from reverse-transcribed RNA isolated from PBMCs of infected animals 93T and 94T were sequenced using Sanger chemistry and sequences combined to yield a consensus SIV genome. This sequence was deposited in NCBI GenBank, accession number PP236443. After filtering as well as counting the UMI and cell barcode by the Cell Ranger count pipeline, the sequenced samples from the same animals were aggregated together to generate a single file containing feature barcode matrices for the downstream analyses. The counting summary statistics generated by 10x Genomics for each sample are shown in **Table S6**. The downstream analyses were then implemented with R (version: 4.3).

### Cell clustering and differentially expressed genes (DEGs)

To cluster the cell and find DEGs for each cell cluster, the feature barcode matrices were analyzed with Seurat R package(^113^) (version: 4.3.0). We removed sparsely expressed genes and low-quality cells and kept genes which had expression in at least 10 cells and the cells with UMI count from 400 to 20,000, gene count from 400 to 10,000, and mitochondrial percentage less than 15%. The final cell counts are shown in **Table S6**. Then the scRNA-seq datasets from six animals were merged to generate a single Seurat Object for further analyses. The normalization, scaling and finding variable genes were performed by SCTransform v2 (SCT).(^114^) After normalization, we performed principal component analyses (PCAs) with a default setting of 50 principal components (PCs) for reducing dimensionality. To minimize the batch effect and integrate the datasets, we implemented harmony(^115^) (version: 1.0) before clustering. The integrated dataset was then subjected to graph-based clustering which used the first 30 PCs and 0.2 as resolution. We selected the top 30 PCs which explain approximately 80% of the total variation. The Uniform Manifold Approximation and Projection (UMAP) was used as a non-linear dimensional reduction method to further visualize the cell clusters. The settings for running UMAP mostly followed the default except for defining the dimensionalities using the first 30 batch effect corrected PCs.

We then characterized each cluster in two ways. First, we examined the expression of general cell markers for microglia (P2RY12, GPR34, and CX3CR1), CNS-associated macrophages (MHC class II), T/NK cells (CD3D, GZMB, and NKG7), B cells (EBF1 and MS4A1), and endothelial cells (RGS5, CLDN5, and ATP1A2) in each cluster. Next, we found the DEGs for each cell cluster by performing the Wilcoxon rank sum test embedded in FindAllMarkers function of Seurat to the data that was normalized by SCT. The positive markers with 0.25-fold change (log-scale) on average that were detected in a minimum of 25% of cells in either of two populations were calculated as biomarkers for each cluster **(Table S2)**. The DEGs that were found for the same macrophage/microglia cluster but under different infection conditions were also identified by using the above test method and criteria **(Table S4)**. When we used the same way to find DEGs between infected cells and uninfected cells (Figure S2 and Table S3), to avoid losing the signal for SIV, we performed FindAllMarkers fucntion on log-normalized data, which was obtained by applying LogNormalize function in Seurat with scale factor as 10000. The average gene expression for the macrophages/microglia in the same cluster was calculated for the uninfected cells and acute-infected cells separately, and then the values were added 1 and converted to log_10_ value for plotting. Given our goal in this study to analyze the myeloid cells in the brain, we then further subsetted the microglial and macrophage clusters and reran the FindAllMarkers function with the aforementioned settings within them separately. The DEGs that were found in subclusters of microglia or macrophages were further used in Gene Ontology (GO) analyses for further characterization.

#### Gene Ontology (GO) and Kyoto Encyclopedia of Genes and Genomes (KEGG) Over-representation analyses (ORA)

Over-representation analysis (ORA) is a widely used approach to determine whether known biological functions or processes are enriched in an experimentally derived gene list (e.g., DEGs), and the p-value in this analysis is calculated by hypergeometric distribution.(^116^) The GO-ORA and KEGG-ORA were all implemented with the clusterProfiler R package(^117^) (version: 4.0.2). The GO analysis uses Entrez Gene identifiers instead of the common gene symbol. Therefore, we converted the common gene symbol to their Entrez Gene identifiers by genome wide annotation package for rhesus macaque (version: 3.18). This package mapped the gene symbol to Entrez Gene identifiers based on NCBI databases (updated on Sep-11, 2023). The featured pathways of each microglial or macrophage cluster were then identified using GO-ORA based on the DEGs, and the biological process (BP) was chosen as subontology for analysis. The KEGG-ORA used the DEGs that were found between different conditions as input to find the upregulated pathways for microglial or macrophage clusters in response to acute SIV infection. The gene annotation of rhesus macaque for KEGG analyses was found in KEGG database (Mmul10, RefSeq). For both GO-ORA and KEGG-ORA, the cutoff for p-value was set to <0.05. The results were visualized by dot plots, bar plots, tree plots, and gene-concept network, which were plotted using enrichplot package (version 1.22.0). All GO-ORA and KEGG-ORA pathways detected with p-value less than 0.05 were summarized in **Table S5**. In the table, the analysis results provided geneRatio and BgRatio, which are the ratio of input genes annotated in a term and the ratio of all genes that are annotated in this term respectively.

#### Trajectory analysis

The monocle3 R package(^52, 118, 119^) (version: 1.2.7) was used to estimate lineage differentiation within the macrophage clusters. We extracted the macrophage clusters from the merged Seurat Object and further constructed single-cell trajectories. The trajectory graph was inferred and fitted to the cell clusters generated by Seurat. The Macro-8 cluster was defined as the root node based on the prior knowledge(^41^) for ordering all macrophages in their pseudotime. For visualization, the UMAP embeddings from Seurat Object was used, the nodes and branches were delineated based on the trajectory analysis, and the cells were colored by their pseudotime. To better compare the pseudotime of cells in Macro-6 and Macro-8, they were further illustrated in box plot using ggplot2 (version: 3.4.4)

### Statistics

Alpha less than 0.05 was considered as a significant difference in all comparisons. The DEGs were found using non-parametric Wilcox rank sum test, and p-values were adjusted based on Bonferroni correction. In ORA analyses, the enrichment p-value is calculated using hypergeometric distribution, and p-value was adjusted in GO-ORA analysis to compare multiple microglial clusters.

## ACKNOWLEDGEMENTS

This work was supported by National Institutes of Health (NIH) grants U01DA053624, R21MH128057, and P30MH062261. We thank Drs. Shilpa Buch and Siddappa Byrareddy, Mr. Moses Apostol, Ms. Brenda Morsey (deceased), as well as all members of our non-human primate/SIV collaborative team at UNMC. We also want to acknowledge UMNC Genomics Core Facility and Flow Cytometry Research Facility for the excellent technical support of scRNA-seq and FACS used in this study, the University of Nebraska Lincoln (UNL) Computing Center for high-performance supercomputer access, the Quantitative Molecular Diagnostics Core of the AIDS, the Cancer Virus Program of the Frederick National Laboratory for expert assistance with viral load measurements, and the Virus Characterization, Isolation and Production Core at the Tulane National Primate Research Center for the SIV_mac_251 stock. The UNMC Genomics Core Facility receives partial support from the National Institute for General Medical Science (NIGMS) INBRE grant P20GM103427, as well as the National Cancer Institute (NCI) Fred and Pamela Buffett Cancer Center support grant P30CA036727. The UNMC Flow Cytometry Research Facility is supported by state funds from the Nebraska Research Initiative and the NCI Fred and Pamela Buffett Cancer Center Support Grant. The UNL Holland Computing Center receives support from the UNL Office of Research and Economic Development and the Nebraska Research Initiative. The Quantitative Molecular Diagnostics Core of the AIDS and Cancer Virus Program of the Frederick National Laboratory is supported in part in part with federal funds from the NCI, NIH, under Contract 75N91019D00024/HHSN261201500003I. The Virus Characterization, Isolation and Production Core at Tulane National Primate Research Center is supported by NIH grant P51OD011104 Major instrumentation at UNMC has been provided by the UNMC Office of the Vice Chancellor for Research, The University of Nebraska Foundation, the Nebraska Banker’s Fund, and by the NIH Shared Instrument Program. This publications’ contents are the sole responsibility of the authors and do not necessarily represent the official views or policies of the NIH or Department of Health and Human Services, nor does mention of trade names, commercial products, or organizations imply endorsement by the U.S. Government.

## Notes

### Competing Interest Statement

The authors have declared no competing interest.

## REFERENCES

1. Threats M, Brawner BM, Montgomery TM, Abrams J, Jemmott LS, Crouch PC, et al. A Review of Recent HIV Prevention Interventions and Future Considerations for Nursing Science. J Assoc Nurses AIDS Care. 2021;32(3):373–91.

2. Burudi EME, Fox HS. Simian immunodeficiency virus model of HIV induced central nervous system dysfunction. Advances in Virus Research. 56: Academic Press; 2001. p. 435-68.

3. Fox HS, Gold LH, Henriksen SJ, Bloom FE. Simian Immunodeficiency Virus: A Model for NeuroAIDS. Neurobiology of Disease. 1997;4(3):265–74.

4. Zink MC, Spelman JP, Robinson RB, Clements JE. SIV infection of macaques--modeling the progression to AIDS dementia. J Neurovirol. 1998;4(3):249–59.

5. Avalos CR, Abreu CM, Queen SE, Li M, Price S, Shirk EN, et al. Brain Macrophages in Simian Immunodeficiency Virus-Infected, Antiretroviral-Suppressed Macaques: a Functional Latent Reservoir. mBio. 2017;8(4).

6. Pantaleo G, Menzo S, Vaccarezza M, Graziosi C, Cohen OJ, Demarest JF, et al. Studies in subjects with long-term nonprogressive human immunodeficiency virus infection. N Engl J Med. 1995;332(4):209–16.

7. Cohen MS, Shaw GM, McMichael AJ, Haynes BF. Acute HIV-1 Infection. N Engl J Med. 2011;364(20):1943–54.

8. Hladik F, Sakchalathorn P, Ballweber L, Lentz G, Fialkow M, Eschenbach D, et al. Initial events in establishing vaginal entry and infection by human immunodeficiency virus type-1. Immunity. 2007;26(2):257–70.

9. Veazey RS, DeMaria M, Chalifoux LV, Shvetz DE, Pauley DR, Knight HL, et al. Gastrointestinal Tract as a Major Site of CD4+ T Cell Depletion and Viral Replication in SIV Infection. Science. 1998;280(5362):427–31.

10. Mattapallil JJ, Douek DC, Hill B, Nishimura Y, Martin M, Roederer M. Massive infection and loss of memory CD4+ T cells in multiple tissues during acute SIV infection. Nature. 2005;434(7037):1093–7.

11. Witwer KW, Gama L, Li M, Bartizal CM, Queen SE, Varrone JJ, et al. Coordinated regulation of SIV replication and immune responses in the CNS. PLoS One. 2009;4(12):e8129.

12. Valcour V, Chalermchai T, Sailasuta N, Marovich M, Lerdlum S, Suttichom D, et al. Central nervous system viral invasion and inflammation during acute HIV infection. J Infect Dis. 2012;206(2):275–82.

13. Clements JE, Babas T, Mankowski JL, Suryanarayana K, Piatak M, Jr., Tarwater PM, et al. The central nervous system as a reservoir for simian immunodeficiency virus (SIV): steady-state levels of SIV DNA in brain from acute through asymptomatic infection. J Infect Dis. 2002;186(7):905–13.

14. Wallet C, De Rovere M, Van Assche J, Daouad F, De Wit S, Gautier V, et al. Microglial Cells: The Main HIV-1 Reservoir in the Brain. Front Cell Infect Microbiol. 2019;9:362.

15. Gannon P, Khan MZ, Kolson DL. Current understanding of HIV-associated neurocognitive disorders pathogenesis. Curr Opin Neurol. 2011;24(3):275–83.

16. Sacktor N, McDermott MP, Marder K, Schifitto G, Selnes OA, McArthur JC, et al. HIV-associated cognitive impairment before and after the advent of combination therapy. J Neurovirol. 2002;8(2):136–42.

17. Kraft-Terry SD, Stothert AR, Buch S, Gendelman HE. HIV-1 neuroimmunity in the era of antiretroviral therapy. Neurobiol Dis. 2010;37(3):542–8.

18. Li Q, Barres BA. Microglia and macrophages in brain homeostasis and disease. Nature Reviews Immunology. 2018;18(4):225–42.

19. Bai R, Song C, Lv S, Chang L, Hua W, Weng W, et al. Role of microglia in HIV-1 infection. AIDS Research and Therapy. 2023;20(1):16.

20. Clay CC, Rodrigues DS, Ho YS, Fallert BA, Janatpour K, Reinhart TA, et al. Neuroinvasion of fluorescein-positive monocytes in acute simian immunodeficiency virus infection. J Virol. 2007;81(21):12040–8.

21. Nowlin BT, Burdo TH, Midkiff CC, Salemi M, Alvarez X, Williams KC. SIV encephalitis lesions are composed of CD163(+) macrophages present in the central nervous system during early SIV infection and SIV-positive macrophages recruited terminally with AIDS. Am J Pathol. 2015;185(6):1649–65.

22. Campbell JH, Ratai E-M, Autissier P, Nolan DJ, Tse S, Miller AD, et al. Anti-α4 Antibody Treatment Blocks Virus Traffic to the Brain and Gut Early, and Stabilizes CNS Injury Late in Infection. PLOS Pathogens. 2014;10(12):e1004533.

23. Thompson KA, Cherry CL, Bell JE, McLean CA. Brain cell reservoirs of latent virus in presymptomatic HIV-infected individuals. Am J Pathol. 2011;179(4):1623–9.

24. Borrajo A, Spuch C, Penedo MA, Olivares JM, Agís-Balboa RC. Important role of microglia in HIV-1 associated neurocognitive disorders and the molecular pathways implicated in its pathogenesis. Ann Med. 2021;53(1):43–69.

25. Castellano P, Prevedel L, Eugenin EA. HIV-infected macrophages and microglia that survive acute infection become viral reservoirs by a mechanism involving Bim. Scientific Reports. 2017;7(1):12866.

26. Chandrasekar AP, Cummins NW, Badley AD. The Role of the BCL-2 Family of Proteins in HIV-1 Pathogenesis and Persistence. Clin Microbiol Rev. 2019;33(1).

27. Hayes GM, Woodroofe MN, Cuzner ML. Microglia are the major cell type expressing MHC class II in human white matter. Journal of the Neurological Sciences. 1987;80(1):25–37.

28. Perlmutter LS, Scott SA, Barrón E, Chui HC. MHC class II-positive microglia in human brain: Association with alzheimer lesions. Journal of Neuroscience Research. 1992;33(4):549–58.

29. Sheffield LG, Berman NEJ. Microglial Expression of MHC Class II Increases in Normal Aging of Nonhuman Primates. Neurobiology of Aging. 1998;19(1):47–55.

30. Zöller T, Schneider A, Kleimeyer C, Masuda T, Potru PS, Pfeifer D, et al. Silencing of TGFβ signalling in microglia results in impaired homeostasis. Nature Communications. 2018;9(1):4011.

31. Sandler NG, Bosinger SE, Estes JD, Zhu RT, Tharp GK, Boritz E, et al. Type I interferon responses in rhesus macaques prevent SIV infection and slow disease progression. Nature. 2014;511(7511):601-5.

32. Vanderford TH, Slichter C, Rogers KA, Lawson BO, Obaede R, Else J, et al. Treatment of SIV-infected sooty mangabeys with a type-I IFN agonist results in decreased virus replication without inducing hyperimmune activation. Blood. 2012;119(24):5750–7.

33. Jacquelin B, Petitjean G, Kunkel D, Liovat AS, Jochems SP, Rogers KA, et al. Innate immune responses and rapid control of inflammation in African green monkeys treated or not with interferon-alpha during primary SIVagm infection. PLoS Pathog. 2014;10(7):e1004241.

34. Parhizkar S, Holtzman DM. APOE mediated neuroinflammation and neurodegeneration in Alzheimer’s disease. Semin Immunol. 2022;59:101594.

35. Jordan CA, Watkins BA, Kufta C, Dubois-Dalcq M. Infection of brain microglial cells by human immunodeficiency virus type 1 is CD4 dependent. J Virol. 1991;65(2):736–42.

36. Lindwasser OW, Chaudhuri R, Bonifacino JS. Mechanisms of CD4 downregulation by the Nef and Vpu proteins of primate immunodeficiency viruses. Curr Mol Med. 2007;7(2):171–84.

37. Beignon AS, McKenna K, Skoberne M, Manches O, DaSilva I, Kavanagh DG, et al. Endocytosis of HIV-1 activates plasmacytoid dendritic cells via Toll-like receptor-viral RNA interactions. J Clin Invest. 2005;115(11):3265–75.

38. Luban J. Innate immune sensing of HIV-1 by dendritic cells. Cell Host Microbe. 2012;12(4):408–18.

39. Iwasaki A. Innate immune recognition of HIV-1. Immunity. 2012;37(3):389–98.

40. Cros J, Cagnard N, Woollard K, Patey N, Zhang SY, Senechal B, et al. Human CD14dim monocytes patrol and sense nucleic acids and viruses via TLR7 and TLR8 receptors. Immunity. 2010;33(3):375–86.

41. Yona S, Kim KW, Wolf Y, Mildner A, Varol D, Breker M, et al. Fate mapping reveals origins and dynamics of monocytes and tissue macrophages under homeostasis. Immunity. 2013;38(1):79–91.

42. Veldhoen M, Hocking RJ, Atkins CJ, Locksley RM, Stockinger B. TGFβ in the context of an inflammatory cytokine milieu supports de novo differentiation of IL-17-producing T cells. Immunity. 2006;24(2):179–89.

43. Tesmer LA, Lundy SK, Sarkar S, Fox DA. Th17 cells in human disease. Immunol Rev. 2008;223:87–113.

44. Gopalakrishnan RM, Aid M, Mercado NB, Davis C, Malik S, Geiger E, et al. Increased IL-6 expression precedes reliable viral detection in the rhesus macaque brain during acute SIV infection. JCI Insight. 2021;6(20).

45. Cummins NW, Sainski-Nguyen AM, Natesampillai S, Aboulnasr F, Kaufmann S, Badley AD. Maintenance of the HIV Reservoir Is Antagonized by Selective BCL2 Inhibition. J Virol. 2017;91(11).

46. Cummins NW, Sainski AM, Dai H, Natesampillai S, Pang YP, Bren GD, et al. Prime, Shock, and Kill: Priming CD4 T Cells from HIV Patients with a BCL-2 Antagonist before HIV Reactivation Reduces HIV Reservoir Size. J Virol. 2016;90(8):4032–48.

47. Prinz M, Masuda T, Wheeler MA, Quintana FJ. Microglia and Central Nervous System-Associated Macrophages-From Origin to Disease Modulation. Annu Rev Immunol. 2021;39:251–77.

48. Wendimu MY, Hooks SB. Microglia Phenotypes in Aging and Neurodegenerative Diseases. Cells. 2022;11(13).

49. Fox HS, Niu M, Morsey BM, Lamberty BG, Emanuel K, Periyasamy P, et al. Morphine suppresses peripheral responses and transforms brain myeloid gene expression to favor neuropathogenesis in SIV infection. Front Immunol. 2022;13:1012884.

50. Trease AJ, Niu M, Morsey B, Guda C, Byrareddy SN, Buch S, et al. Antiretroviral therapy restores the homeostatic state of microglia in SIV-infected rhesus macaques. J Leukoc Biol. 2022;112(5):969–81.

51. Pietrowski MJ, Gabr AA, Kozlov S, Blum D, Halle A, Carvalho K. Glial Purinergic Signaling in Neurodegeneration. Front Neurol. 2021;12:654850.

52. Grubman A, Chew G, Ouyang JF, Sun G, Choo XY, McLean C, et al. A single-cell atlas of entorhinal cortex from individuals with Alzheimer’s disease reveals cell-type-specific gene expression regulation. Nature Neuroscience. 2019;22(12):2087–97.

53. Ruganzu JB, Peng X, He Y, Wu X, Zheng Q, Ding B, et al. Downregulation of TREM2 expression exacerbates neuroinflammatory responses through TLR4-mediated MAPK signaling pathway in a transgenic mouse model of Alzheimer’s disease. Mol Immunol. 2022;142:22–36.

54. Cardona AE, Pioro EP, Sasse ME, Kostenko V, Cardona SM, Dijkstra IM, et al. Control of microglial neurotoxicity by the fractalkine receptor. Nature Neuroscience. 2006;9(7):917–24.

55. Pettas S, Karagianni K, Kanata E, Chatziefstathiou A, Christoudia N, Xanthopoulos K, et al. Profiling Microglia through Single-Cell RNA Sequencing over the Course of Development, Aging, and Disease. Cells. 2022;11(15).

56. Buttgereit A, Lelios I, Yu X, Vrohlings M, Krakoski NR, Gautier EL, et al. Sall1 is a transcriptional regulator defining microglia identity and function. Nature Immunology. 2016;17(12):1397–406.

57. Butovsky O, Jedrychowski MP, Moore CS, Cialic R, Lanser AJ, Gabriely G, et al. Identification of a unique TGF-β-dependent molecular and functional signature in microglia. Nat Neurosci. 2014;17(1):131–43.

58. Gosselin D, Link VM, Romanoski CE, Fonseca GJ, Eichenfield DZ, Spann NJ, et al. Environment drives selection and function of enhancers controlling tissue-specific macrophage identities. Cell. 2014;159(6):1327–40.

59. Spittau B, Dokalis N, Prinz M. The Role of TGFβ Signaling in Microglia Maturation and Activation. Trends in Immunology. 2020;41(9):836–48.

60. Sankowski R, Böttcher C, Masuda T, Geirsdottir L, Sagar, Sindram E, et al. Mapping microglia states in the human brain through the integration of high-dimensional techniques. Nature Neuroscience. 2019;22(12):2098–110.

61. Geirsdottir L, David E, Keren-Shaul H, Weiner A, Bohlen SC, Neuber J, et al. Cross-Species Single-Cell Analysis Reveals Divergence of the Primate Microglia Program. Cell. 2019;179(7):1609–22.e16.

62. Masuda T, Sankowski R, Staszewski O, Böttcher C, Amann L, Sagar, et al. Spatial and temporal heterogeneity of mouse and human microglia at single-cell resolution. Nature. 2019;566(7744):388-92.

63. Croft NP, Smith SA, Pickering J, Sidney J, Peters B, Faridi P, et al. Most viral peptides displayed by class I MHC on infected cells are immunogenic. Proceedings of the National Academy of Sciences. 2019;116(8):3112–7.

64. Barber SA, Herbst DS, Bullock BT, Gama L, Clements JE. Innate immune responses and control of acute simian immunodeficiency virus replication in the central nervous system. Journal of NeuroVirology. 2004;10(1):15–20.

65. Alammar L, Gama L, Clements JE. Simian immunodeficiency virus infection in the brain and lung leads to differential type I IFN signaling during acute infection. J Immunol. 2011;186(7):4008–18.

66. Bosinger SE, Li Q, Gordon SN, Klatt NR, Duan L, Xu L, et al. Global genomic analysis reveals rapid control of a robust innate response in SIV-infected sooty mangabeys. J Clin Invest. 2009;119(12):3556–72.

67. Echebli N, Tchitchek N, Dupuy S, Bruel T, Peireira Bittencourt Passaes C, Bosquet N, et al. Stage-specific IFN-induced and IFN gene expression reveal convergence of type I and type II IFN and highlight their role in both acute and chronic stage of pathogenic SIV infection. PLoS One. 2018;13(1):e0190334.

68. Roberts ES, Burudi EM, Flynn C, Madden LJ, Roinick KL, Watry DD, et al. Acute SIV infection of the brain leads to upregulation of IL6 and interferon-regulated genes: expression patterns throughout disease progression and impact on neuroAIDS. J Neuroimmunol. 2004;157(1-2):81–92.

69. Hernández JC, Stevenson M, Latz E, Urcuqui-Inchima S. HIV type 1 infection up-regulates TLR2 and TLR4 expression and function in vivo and in vitro. AIDS Res Hum Retroviruses. 2012;28(10):1313–28.

70. Del Cornò M, Cappon A, Donninelli G, Varano B, Marra F, Gessani S. HIV-1 gp120 signaling through TLR4 modulates innate immune activation in human macrophages and the biology of hepatic stellate cells. J Leukoc Biol. 2016;100(3):599–606.

71. Mandl JN, Barry AP, Vanderford TH, Kozyr N, Chavan R, Klucking S, et al. Divergent TLR7 and TLR9 signaling and type I interferon production distinguish pathogenic and nonpathogenic AIDS virus infections. Nat Med. 2008;14(10):1077–87.

72. Henrick BM, Nag K, Yao XD, Drannik AG, Aldrovandi GM, Rosenthal KL. Milk matters: soluble Toll-like receptor 2 (sTLR2) in breast milk significantly inhibits HIV-1 infection and inflammation. PLoS One. 2012;7(7):e40138.

73. Henrick BM, Yao XD, Drannik AG, Abimiku A, Rosenthal KL. Soluble toll-like receptor 2 is significantly elevated in HIV-1 infected breast milk and inhibits HIV-1 induced cellular activation, inflammation and infection. Aids. 2014;28(14):2023–32.

74. Henrick BM, Yao XD, Rosenthal KL. HIV-1 Structural Proteins Serve as PAMPs for TLR2 Heterodimers Significantly Increasing Infection and Innate Immune Activation. Front Immunol. 2015;6:426.

75. Duverger A, Wolschendorf F, Zhang M, Wagner F, Hatcher B, Jones J, et al. An AP-1 binding site in the enhancer/core element of the HIV-1 promoter controls the ability of HIV-1 to establish latent infection. J Virol. 2013;87(4):2264–77.

76. Hoshino S, Konishi M, Mori M, Shimura M, Nishitani C, Kuroki Y, et al. HIV-1 Vpr induces TLR4/MyD88-mediated IL-6 production and reactivates viral production from latency. Journal of Leukocyte Biology. 2010;87(6):1133–43.

77. Varin A, Decrion AZ, Sabbah E, Quivy V, Sire J, Van Lint C, et al. Synthetic Vpr protein activates activator protein-1, c-Jun N-terminal kinase, and NF-kappaB and stimulates HIV-1 transcription in promonocytic cells and primary macrophages. J Biol Chem. 2005;280(52):42557–67.

78. Liu R, Lin Y, Jia R, Geng Y, Liang C, Tan J, et al. HIV-1 Vpr stimulates NF-κB and AP-1 signaling by activating TAK1. Retrovirology. 2014;11(1):45.

79. Nixon CC, Mavigner M, Sampey GC, Brooks AD, Spagnuolo RA, Irlbeck DM, et al. Systemic HIV and SIV latency reversal via non-canonical NF-κB signalling in vivo. Nature. 2020;578(7793):160-5.

80. Goldmann T, Wieghofer P, Jordão MJ, Prutek F, Hagemeyer N, Frenzel K, et al. Origin, fate and dynamics of macrophages at central nervous system interfaces. Nat Immunol. 2016;17(7):797–805.

81. Prinz M, Erny D, Hagemeyer N. Ontogeny and homeostasis of CNS myeloid cells. Nature Immunology. 2017;18(4):385–92.

82. Nowlin BT, Wang J, Schafer JL, Autissier P, Burdo TH, Williams KC. Monocyte subsets exhibit transcriptional plasticity and a shared response to interferon in SIV-infected rhesus macaques. Journal of Leukocyte Biology. 2018;103(1):141–55.

83. Ronning KE, Karlen SJ, Burns ME. Structural and functional distinctions of co-resident microglia and monocyte-derived macrophages after retinal degeneration. Journal of Neuroinflammation. 2022;19(1):299.

84. Eales LJ, Farrant J, Helbert M, Pinching AJ. Peripheral blood dendritic cells in persons with AIDS and AIDS related complex: loss of high intensity class II antigen expression and function. Clin Exp Immunol. 1988;71(3):423–7.

85. Polyak S, Chen H, Hirsch D, George I, Hershberg R, Sperber K. Impaired class II expression and antigen uptake in monocytic cells after HIV-1 infection. J Immunol. 1997;159(5):2177–88.

86. Kanazawa S, Okamoto T, Peterlin BM. Tat Competes with CIITA for the Binding to P-TEFb and Blocks the Expression of MHC Class II Genes in HIV Infection. Immunity. 2000;12(1):61–70.

87. Stumptner-Cuvelette P, Morchoisne S, Dugast M, Le Gall S, Raposo G, Schwartz O, et al. HIV-1 Nef impairs MHC class II antigen presentation and surface expression. Proceedings of the National Academy of Sciences. 2001;98(21):12144–9.

88. Williams DW, Veenstra M, Gaskill PJ, Morgello S, Calderon TM, Berman JW. Monocytes mediate HIV neuropathogenesis: mechanisms that contribute to HIV associated neurocognitive disorders. Curr HIV Res. 2014;12(2):85–96.

89. Castellano P, Prevedel L, Valdebenito S, Eugenin EA. HIV infection and latency induce a unique metabolic signature in human macrophages. Scientific Reports. 2019;9(1):3941.

90. Saylor D, Dickens AM, Sacktor N, Haughey N, Slusher B, Pletnikov M, et al. HIV-associated neurocognitive disorder--pathogenesis and prospects for treatment. Nat Rev Neurol. 2016;12(4):234–48.

91. Abassi M, Morawski BM, Nakigozi G, Nakasujja N, Kong X, Meya DB, et al. Cerebrospinal fluid biomarkers and HIV-associated neurocognitive disorders in HIV-infected individuals in Rakai, Uganda. J Neurovirol. 2017;23(3):369–75.

92. Jessen Krut J, Mellberg T, Price RW, Hagberg L, Fuchs D, Rosengren L, et al. Biomarker evidence of axonal injury in neuroasymptomatic HIV-1 patients. PLoS One. 2014;9(2):e88591.

93. Borda JT, Alvarez X, Mohan M, Hasegawa A, Bernardino A, Jean S, et al. CD163, a marker of perivascular macrophages, is up-regulated by microglia in simian immunodeficiency virus encephalitis after haptoglobin-hemoglobin complex stimulation and is suggestive of breakdown of the blood-brain barrier. Am J Pathol. 2008;172(3):725–37.

94. Fields JA, Ellis RJ. HIV in the cART era and the mitochondrial: immune interface in the CNS. Int Rev Neurobiol. 2019;145:29–65.

95. Fazeli PL, Crowe M, Ross LA, Wadley V, Ball K, Vance DE. Cognitive Functioning in Adults Aging with HIV: A Cross-Sectional Analysis of Cognitive Subtypes and Influential Factors. J Clin Res HIV AIDS Prev. 2014;1(4):155–69.

96. Valcour V, Shikuma C, Shiramizu B, Watters M, Poff P, Selnes O, et al. Higher frequency of dementia in older HIV-1 individuals: the Hawaii Aging with HIV-1 Cohort. Neurology. 2004;63(5):822–7.

97. Filgueira L, Larionov A, Lannes N. The Influence of Virus Infection on Microglia and Accelerated Brain Aging. Cells [Internet]. 2021; 10(7).

98. Lew BJ, Schantell MD, O’Neill J, Morsey B, Wang T, Ideker T, et al. Reductions in Gray Matter Linked to Epigenetic HIV-Associated Accelerated Aging. Cereb Cortex. 2021;31(8):3752–63.

99. Proskovec AL, Rezich MT, O’Neill J, Morsey B, Wang T, Ideker T, et al. Association of Epigenetic Metrics of Biological Age With Cortical Thickness. JAMA Netw Open. 2020;3(9):e2015428.

100. Gross AM, Jaeger PA, Kreisberg JF, Licon K, Jepsen KL, Khosroheidari M, et al. Methylome-wide Analysis of Chronic HIV Infection Reveals Five-Year Increase in Biological Age and Epigenetic Targeting of HLA. Mol Cell. 2016;62(2):157–68.

101. Horvath S, Levine AJ. HIV-1 Infection Accelerates Age According to the Epigenetic Clock. J Infect Dis. 2015;212(10):1563–73.

102. Shiramizu B, Ananworanich J, Chalermchai T, Siangphoe U, Troelstrup D, Shikuma C, et al. Failure to clear intra-monocyte HIV infection linked to persistent neuropsychological testing impairment after first-line combined antiretroviral therapy. J Neurovirol. 2012;18(1):69–73.

103. Shikuma CM, Nakamoto B, Shiramizu B, Liang CY, DeGruttola V, Bennett K, et al. Antiretroviral monocyte efficacy score linked to cognitive impairment in HIV. Antivir Ther. 2012;17(7):1233–42.

104. Krajewski S, James HJ, Ross J, Blumberg BM, Epstein LG, Gendelman HE, et al. Expression of pro-and anti-apoptosis gene products in brains from paediatric patients with HIV-1 encephalitis. Neuropathology and Applied Neurobiology. 1997;23(3):242–53.

105. Campbell Grant R, To Rachel K, Spector Stephen A. TREM-1 Protects HIV-1-Infected Macrophages from Apoptosis through Maintenance of Mitochondrial Function. mBio. 2019;10(6):10.1128/mbio.02638-19.

106. Sanchez-Moral L, Ràfols N, Martori C, Paul T, Téllez É, Sarrias MR. Multifaceted Roles of CD5L in Infectious and Sterile Inflammation. Int J Mol Sci. 2021;22(8).

107. Miyazaki T, Hirokami Y, Matsuhashi N, Takatsuka H, Naito M. Increased susceptibility of thymocytes to apoptosis in mice lacking AIM, a novel murine macrophage-derived soluble factor belonging to the scavenger receptor cysteine-rich domain superfamily. J Exp Med. 1999;189(2):413–22.

108. Wishart CL, Spiteri AG, Locatelli G, King NJC. Integrating transcriptomic datasets across neurological disease identifies unique myeloid subpopulations driving disease-specific signatures. Glia. 2023;71(4):904–25.

109. Hansen SG, Piatak M, Ventura AB, Hughes CM, Gilbride RM, Ford JC, et al. Addendum: Immune clearance of highly pathogenic SIV infection. Nature. 2017;547(7661):123–4.

110. Hansen SG, Piatak M, Jr., Ventura AB, Hughes CM, Gilbride RM, Ford JC, et al. Immune clearance of highly pathogenic SIV infection. Nature. 2013;502(7469):100–4.

111. Venneti S, Bonneh-Barkay D, Lopresti BJ, Bissel SJ, Wang G, Mathis CA, et al. Longitudinal in vivo positron emission tomography imaging of infected and activated brain macrophages in a macaque model of human immunodeficiency virus encephalitis correlates with central and peripheral markers of encephalitis and areas of synaptic degeneration. Am J Pathol. 2008;172(6):1603–16.

112. Morsey B, Niu M, Dyavar SR, Fletcher CV, Lamberty BG, Emanuel K, et al. Cryopreservation of microglia enables single-cell RNA sequencing with minimal effects on disease-related gene expression patterns. iScience. 2021;24(4):102357.

113. Hao Y, Hao S, Andersen-Nissen E, Mauck WM, 3rd, Zheng S, Butler A, et al. Integrated analysis of multimodal single-cell data. Cell. 2021;184(13):3573–87.e29.

114. Choudhary S, Satija R. Comparison and evaluation of statistical error models for scRNA-seq. Genome Biology. 2022;23(1):27.

115. Korsunsky I, Millard N, Fan J, Slowikowski K, Zhang F, Wei K, et al. Fast, sensitive and accurate integration of single-cell data with Harmony. Nature Methods. 2019;16(12):1289–96.

116. Boyle EI, Weng S, Gollub J, Jin H, Botstein D, Cherry JM, et al. GO::TermFinder--open source software for accessing Gene Ontology information and finding significantly enriched Gene Ontology terms associated with a list of genes. Bioinformatics. 2004;20(18):3710–5.

117. Wu T, Hu E, Xu S, Chen M, Guo P, Dai Z, et al. clusterProfiler 4.0: A universal enrichment tool for interpreting omics data. Innovation (Camb). 2021;2(3):100141.

118. Qiu X, Mao Q, Tang Y, Wang L, Chawla R, Pliner HA, et al. Reversed graph embedding resolves complex single-cell trajectories. Nature Methods. 2017;14(10):979–82.

119. Trapnell C, Cacchiarelli D, Grimsby J, Pokharel P, Li S, Morse M, et al. The dynamics and regulators of cell fate decisions are revealed by pseudotemporal ordering of single cells. Nature Biotechnology. 2014;32(4):381–6.

